# Radiation-Induced Autophagy Regulates the Fibroblast Mitochondrial Stress Response and Crosstalk with Triple-Negative Breast Cancer Cells

**DOI:** 10.1101/2022.05.17.492249

**Authors:** Kevin C. Corn, Shannon E. Martello, Vinay K. Menon, Lucy S. Britto, Kara M. Simmons, Youssef K. Mohamed, Yoanna I. Ivanova, Abtin A. Ghelmansaraei, Sara A. Weidenbach, Tian Zhu, Evan Krystofiak, Jamey D. Young, Vivian Gama, Marjan Rafat

## Abstract

Patients with triple-negative breast cancer (TNBC) experience high recurrence rates despite current interventions, which includes radiation therapy (RT). Tumor cells thought to be involved in recurrence survive in part due to their interactions with irradiated fibroblasts following treatment. How fibroblasts metabolically respond to RT and influence the behavior of TNBC cells is poorly understood. In this study, we demonstrate that irradiated fibroblasts undergo a mitochondrial stress response that is regulated by autophagy, resulting in a metabolic profile characterized by high levels of mitochondrial respiration and fatty acid oxidation. This stress response in fibroblasts induces an aggressive phenotype in TNBC cells that is mitigated when fibroblast autophagy is blocked. Our work reveals how a metabolic stress response in irradiated fibroblasts and crosstalk with TNBC cells leads to a microenvironment conducive to recurrence.

**Graphical Abstract:** 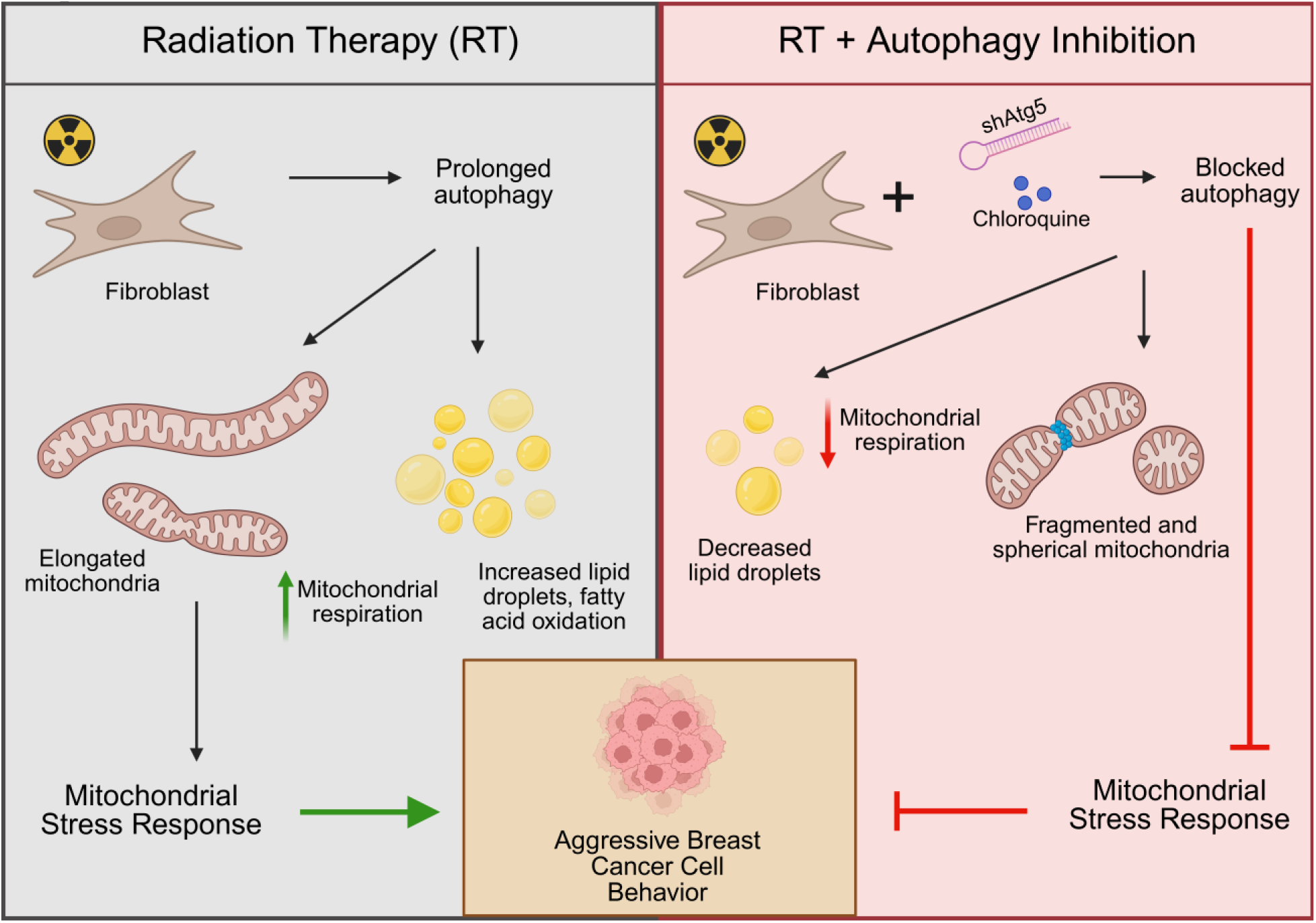

## Introduction

Breast cancer, the most diagnosed cancer worldwide, is one of the deadliest forms of cancer for women^1^. Approximately 15% of breast cancer patients are diagnosed with triple-negative breast cancer (TNBC), a highly aggressive and metastatic subtype that lacks targeted therapies^2,3^. Treatment for TNBC typically consists of chemotherapy, surgery, immune checkpoint inhibition, and radiation therapy (RT)^4–7^. The primary goal of RT is to eliminate residual cancer cells following surgery, and it is largely effective at reducing the incidence of recurrent disease^8,9^. Although improved therapies have led to better outcomes and decreased off-target effects, patients with TNBC still experience high recurrence rates^10–14^. The recruitment of circulating tumor cells (CTCs) to irradiated sites has been identified as a potential contributing factor to recurrence beyond residual tumor cells^15–17^. Therefore, the interactions between tumor cells and the stroma following RT may be important in driving treatment failure and recurrence^18^.

Altered cellular metabolism is a hallmark of cancer progression, and unique metabolic profiles are critical for CTC survival and tumor cell treatment resistance^19,20^. One common metabolic transformation among cancer cell types is a decrease in oxidative phosphorylation and increased reliance on glycolysis, even under aerobic conditions^21^. This is due to both a demand for molecular anabolism and genetic mutations that cause mitochondrial dysfunction^22^. These changes in mitochondrial metabolism can result in increased generation of reactive oxygen species (ROS) leading to genomic instability, further mitochondrial damage, and activated signaling pathways that promote an aggressive phenotype and metastatic behavior^23–25^. Cellular stress responses allow for tumor cell survival after this increased mitochondrial oxidative damage^26–28^, and targeting the survival stress response is increasingly becoming an area of interest in the development of cancer therapeutics^29^. Increased mitochondrial ROS can also result in high autophagy levels to clear damaged organelles, which improves tumor cell survival^30,31^. These stress responses are closely linked to alterations in lipid metabolism, which is known to be critical for tumor cell treatment resistance and survival during the metastatic cascade^32–34^. For example, residual radioresistant cancer cells adopt an altered lipid metabolic profile that may promote recurrence^19,35^, and both increased *de novo* fatty acid synthesis and fatty acid oxidation (FAO) are linked to radioresistance in breast and other cancers treated by RT as well as CTC survival^36–40^.

Fibroblasts are important cellular contributors to the post-RT wound healing response and are subject to off-target effects of RT, which may modulate their interactions with recruited CTCs^41,42^. Determining how tissue fibroblasts metabolically respond to radiation is therefore crucial for understanding recurrence. RT studies typically focus on cancer-associated fibroblasts (CAFs) and their interactions with tumor cells. As normal fibroblasts are activated and transformed into CAFs, they too undergo metabolic reprogramming that can mirror transformations in tumor cells^43^. Mitochondrial DNA damage, mitochondrial dysfunction, and oxidative stress observed in CAFs can influence their crosstalk with tumor cells and promote progression^44–47^. CAFs have also been shown to alter the metabolic profile of cancer cells, both through metabolite transfer and tumor cell metabolic pathway activation^48–52^. How RT modifies the metabolic profile of normal fibroblasts at the treatment site and how these changes affect tumor cell behavior are unknown in TNBC recurrence.

In this study, we show how radiation alters fibroblast metabolism through changes in mitochondrial morphology, autophagic flux, and mitochondrial respiration. We show that irradiated fibroblasts have an elongated mitochondrial morphology that is indicative of a stress response to RT, which is regulated by autophagy. These processes, in addition to increased FAO, increase fibroblast mitochondrial respiration. These highly energetic fibroblasts increase TNBC cell migration rates and tumor spheroid outgrowth, and this phenomenon is reversed when the mitochondrial stress response is disrupted by autophagy inhibition in irradiated fibroblasts. Our study reveals a metabolic mechanism in stromal cells that can be targeted to reduce TNBC aggressiveness and improve the survival of TNBC patients following RT.

## Results

### Radiation causes mitochondrial elongation in fibroblasts

We hypothesized that irradiated fibroblasts have unique metabolic signatures compared to unirradiated fibroblasts. We irradiated murine NIH 3T3 fibroblasts to a total dose of 10 Gy and cultured cells for 7 days. To gain insight into the cellular metabolic changes after radiation, we used transmission electron microscopy (TEM) to visualize organelle morphology (**Fig. 1A**). In unirradiated fibroblasts, mitochondria appeared oval and had few cristae folds within the lumen. Irradiated cells exhibited substantially elongated mitochondria with increased cristae density. We validated the observed mitochondrial elongation through immunofluorescence staining of ATP5A1, a subunit of ATP synthase and mitochondrial marker^53^ (**Fig. 1B**). Quantification of mitochondrial aspect ratio revealed a significant increase in irradiated cells compared to unirradiated cells at both 3 and 7 days after RT (**Fig. 1C–D**). This resulted in an approximate 2-fold increase in overall mitochondrial area per cell by day 7 (**Fig. 1E**), suggesting not only increased fusion but also increased mitochondrial mass following RT. We next evaluated expression of *Mfn1, Mfn2,* and *Opa1*, genes important in regulating mitochondrial fusion^54^, as well as *Sirt5*, which has not only been found to be important in mitochondrial fusion but also mitochondrial respiration and FAO maintenance^55–57^, and found an increasing trend of expression across both early and late time points following RT (**Fig. 1F**). We additionally confirmed these trends in immortalized human reduction mammoplasty fibroblasts (iMFs; **Fig. 1G, S1A**). We also isolated mouse mammary fat pads and irradiated them to a dose of 20 Gy *ex vivo*. This model allowed us to observe the influence of other cells within the breast microenvironment on irradiated fibroblasts without the confounding factor of infiltrating unirradiated cells. *Ex vivo* results confirmed increased mitochondrial staining in tissue fibroblasts (**Fig. 1H**). Taken together, these data suggest metabolic rewiring within irradiated fibroblasts due to changes in mitochondrial morphology.

**Figure 1.**
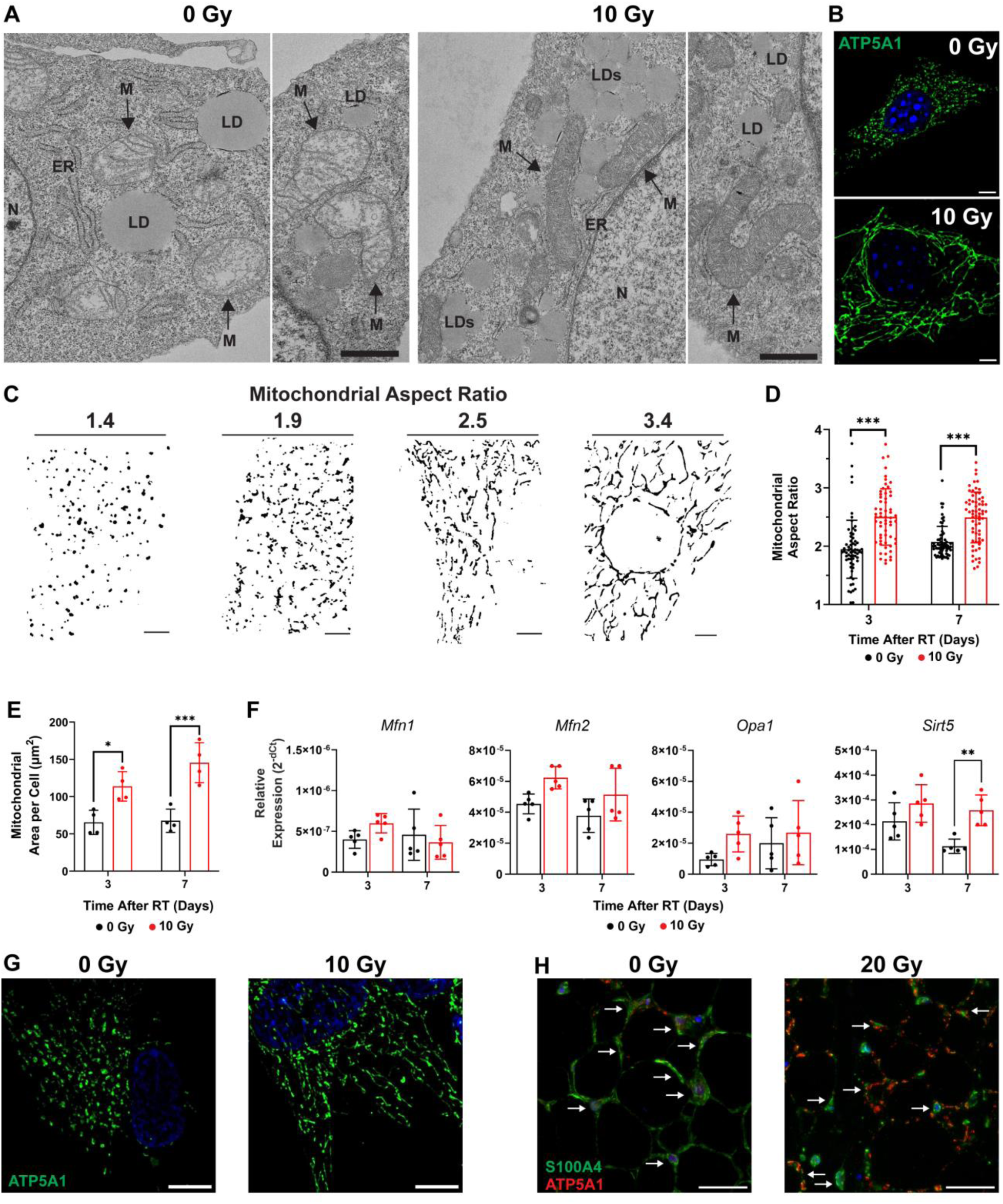
Radiation induces mitochondrial fusion and elongation in fibroblasts. **(A)** Representative transmission electron microscopy (TEM) images of unirradiated (0 Gy) and irradiated (10 Gy) 3T3 fibroblasts 7 days after RT. N=nucleus, ER=endoplasmic reticulum, M=mitochondria, LD=lipid droplet. Arrows also indicate mitochondria. Scale bars are 800 nm. **(B)** Representative immunofluorescence (IF) images of ATP5A1 (green) and nuclear (blue) staining at 7 days after RT. Scale bars are 10 µm. **(C)** Example binary images of mitochondria used for aspect ratio analysis. Images were selected to show how small changes in aspect ratio result in large changes in elongation. Scale bars are 5 µm. **(D)** Quantification of the mitochondrial aspect ratio at 3 and 7 days after RT in unirradiated and irradiated 3T3 fibroblasts. Each data point represents mitochondria from individual cells with n=4 independent replicates. **(E)** Quantification of the total area of ATP5A1 staining per cell in 3- and 7-day post-RT fibroblasts (n=4). **(F)** Relative gene expression of *Mfn1, Mfn2, Opa1,* and *Sirt5* in 3- and 7-day post-RT fibroblasts (n=5). **(G)** Representative images of human immortalized mammary fibroblasts (iMFs) with ATP5A1 (green) and nuclear (blue) staining 7 days after RT. Scale bars are 10 µm. **(H)** Representative images of mouse mammary fat pad cross-sections 7 days after RT to a dose of 20 Gy with S100A4 (green), ATP5A1 (red), and nuclear (blue) staining. White arrows indicate S100A4+ fibroblasts. Scale bars are 30 µm. For **D**, statistical significance was determined by a Mood’s median test for non-equal variances with ***p<0.001. For **E**, statistical analysis was determined by one-way ANOVA with Tukey simultaneous tests for equal variances with *p<0.05, **p<0.01, and ***p<0.001. For **F**, statistical analysis determined by two-way ANOVA with Sidak correction with **p<0.01. Error bars show standard deviation.

### Irradiated fibroblasts maintain high levels of autophagic flux

When analyzing changes in fibroblast morphology through TEM, we also detected more vacuole-like structures in irradiated compared to unirradiated cells (**Fig. 2A**, **S1B**). We determined an increase in lysosomal associated membrane protein 1 (LAMP-1; **Fig. 2B**) expression in irradiated cells, suggesting that a portion of those structures are lysosomes. In tumor cells, autophagy has been demonstrated as a mechanism for the removal of biomolecules and cellular compartments damaged by ROS generated from RT^58,59^, albeit at relatively short time periods after radiation. With the observation of increased lysosomal structures at 7 days after RT in fibroblasts (**Fig. 2B**), we hypothesized that these lysosomes indicate continued autophagy upregulation. To test this hypothesis, we stained for microtubule-associated protein 1A/1B-light chain 3B (LC3B)^60^. Irradiated cells exhibited bright LC3B puncta formation starting 3 days after RT (**Fig. S1C**, **2C**), indicating the formation of autophagosomes. Because the ATG5-ATG12 conjugated complex is involved in the elongation and maturation of the autophagosome membrane^61^, we analyzed ATG5 protein expression through Western blotting. Increased expression was detected in irradiated cells at both 3 and 7 days after RT (**Fig. 2D–E**).

**Figure 2.**
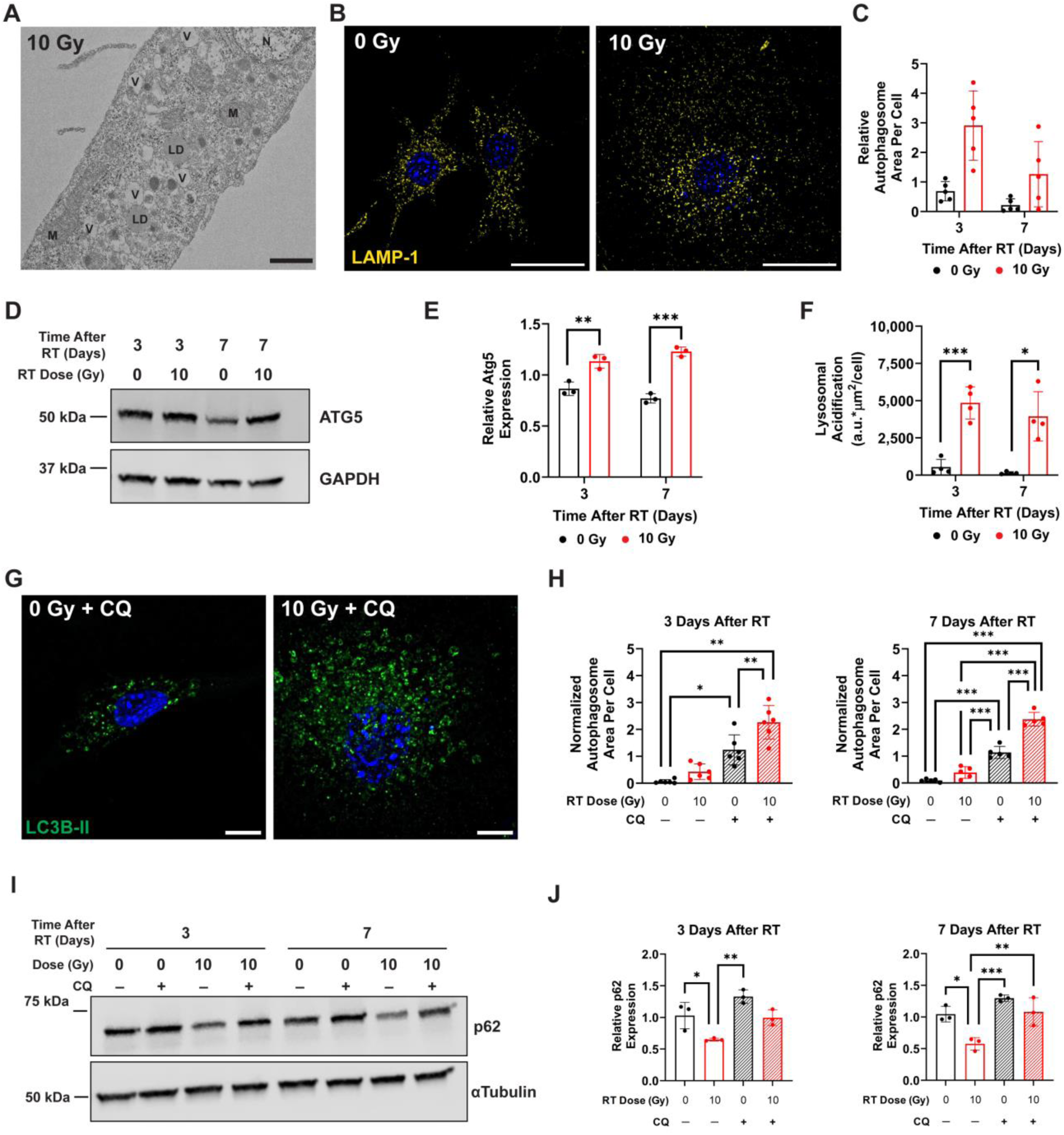
Radiation induces prolonged autophagy upregulation in fibroblasts. **(A)** Representative TEM image of an irradiated 3T3 fibroblast 7 days after RT. V=vacuole-like structure, N=nucleus, M=mitochondria, LD=lipid droplet. Scale bar is 800 nm. **(B)** Representative images of LAMP-1 (yellow) and nuclear (blue) staining of unirradiated and irradiated 3T3 fibroblasts 7 days after RT. Scale bars are 30 µm. **(C)** Quantification of relative autophagosome area per cell, normalized to the biological replicate average, in 3T3 fibroblasts at 3 and 7 days after RT (n=5). **(D)** Representative Western blot for ATG5 with GAPDH as loading control for both irradiated and unirradiated fibroblasts at 3 and 7 days after RT. **(E)** Quantification of relative protein expression of ATG5 normalized to GAPDH loading control expression in irradiated and unirradiated fibroblasts (n=3). **(F)** Quantification of live cell lysosomal staining in irradiated and unirradiated 3T3 fibroblasts (n=4). **(G)** Representative images of LC3B (green) and nuclear (blue) staining in irradiated and unirradiated 3T3s 3 days after RT following autophagy inhibition with chloroquine (CQ). Autophagosomes are identified by the formation of bright LC3B puncta. Scale bars are 10 µm. **(H)** Corresponding quantification of relative autophagosome area per cell, normalized to the biological replicate average, in irradiated and control 3T3s at 3 and 7 days after RT (n=6, 3-day, non-equal variances; n=5, 7-day, equal variances). **(I)** Representative Western blot showing p62 expression with α-tubulin loading control for 3T3s at both 3 and 7 days after RT with and without CQ incubation. **(J)** Quantification of p62 protein expression relative to α-tubulin loading control in irradiated and unirradiated fibroblasts (n=3). For **E**, **F**, **H** (7 day), and **J**, statistical analysis was determined by one-way ANOVA with Tukey simultaneous tests for equal variances with *p<0.05, **p<0.01, and ***p<0.001. For **H** (3 day), statistical analysis was determined by one-way ANOVA with Games-Howell pairwise comparison corrections for non-equal variances with *p<0.05 and **p<0.01. Error bars show standard deviation.

Although increased lysosomal staining and LC3B puncta formation may indicate higher autophagic flux, these results can also be observed in cells with defective autophagy^62^, which has been shown in irradiated cells^63^. To confirm defective autophagy was not the cause of our observations, we analyzed lysosomal acidification through LysoTracker live cell imaging and measured a significant increase in signal (**Fig. S1D**, **2F**). To further evaluate increased autophagic flux, we utilized chloroquine (CQ) to block lysosomal acidification and the fusion of autophagosomes with lysosomes. A dose of 100 µM was sufficient for blocking lysosomal acidification in 3T3 fibroblasts (**Fig. S1E**). CQ incubation increased the formation of LC3B puncta (**Fig. 2G**) compared to untreated controls (**Fig. S1C**), and the highest autophagosome accumulation was observed in the irradiated conditions with CQ incubation (**Fig. 2H**). This suggests that irradiated cells have a higher autophagic flux than unirradiated cells up to 7 days after RT. We additionally analyzed protein expression of Sequestosome-1/p62 with and without CQ incubation. We confirmed a decrease in p62 expression in irradiated fibroblasts and an increase upon CQ incubation (**Fig. 2I–J**), demonstrating upregulated autophagic flux up to 7 days after RT^64^.

### Radiation-induced autophagy promotes mitochondrial elongation

To test the hypothesis that autophagy regulates mitochondrial morphology, we blocked autophagy with CQ and observed an absence of elongated mitochondria in irradiated fibroblasts through TEM (**Fig. 3A**). We again confirmed this morphology difference through IF staining for ATP5A1 (**Fig. 3B**), which showed that the increased mitochondrial aspect ratio post-radiation was almost completely abrogated, especially 7 days after RT once CQ was introduced (**Fig. 3C**). Short-term CQ treatment is widely used to block autophagy through lysosomal deacidification as off-target effects may be severe at longer times^65^. To validate the decrease in mitochondrial elongation due to autophagy inhibition, we developed an ATG5 knockdown 3T3 fibroblast line (shAtg5) to prevent the maturation of autophagosomes and block autophagic flux. ATG5 expression was reduced by 89% in shAtg5 cells compared to scramble controls (shScr, **Fig. 3D**), and autophagic flux as determined by the formation of LC3B-II was successfully decreased after irradiation up to 7-days post-RT (**Fig. 3E**).

**Figure 3.**
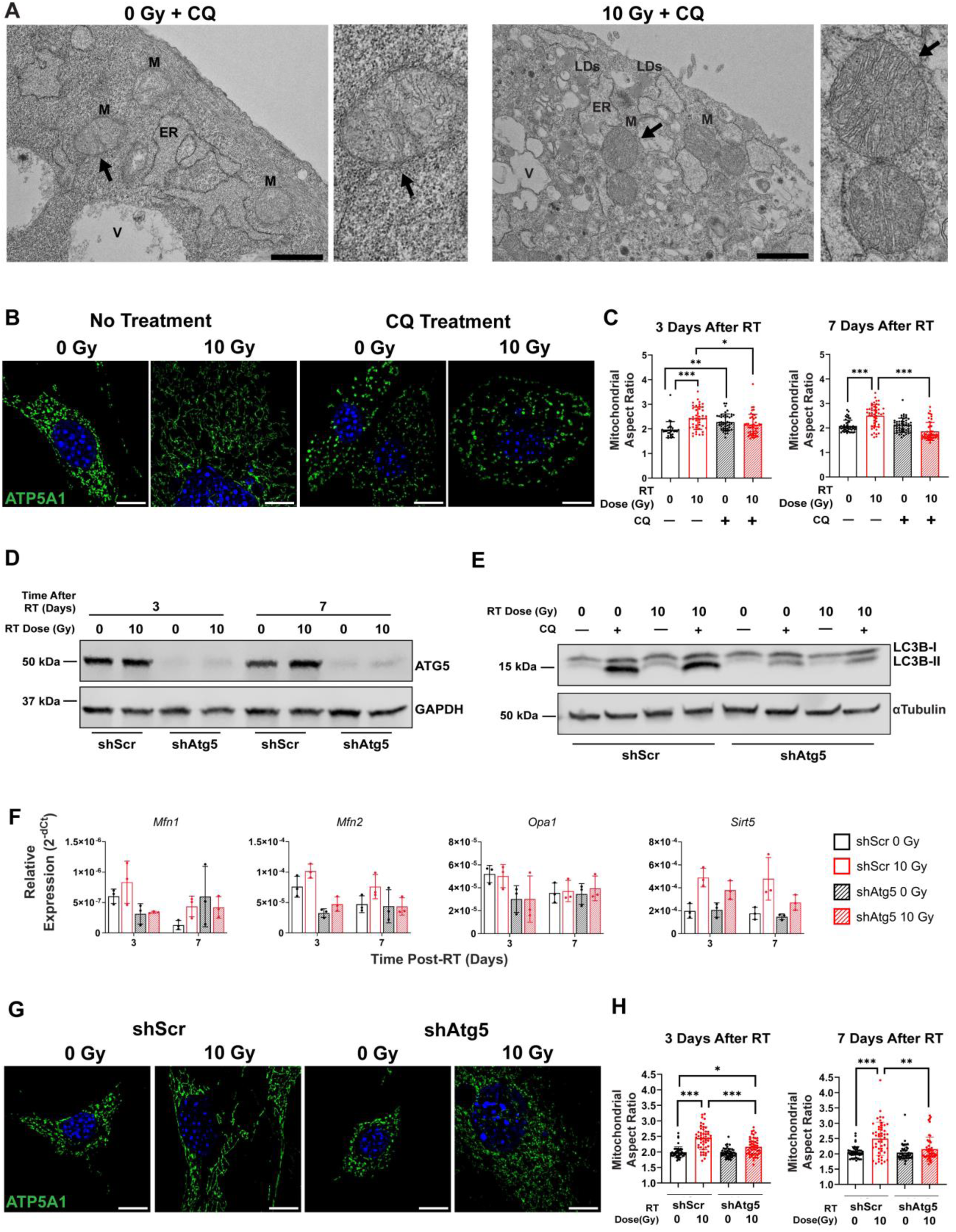
Blocking autophagy in irradiated fibroblasts reverses mitochondrial fusion and elongation. **(A)** Representative transmission electron microscopy (TEM) images of unirradiated and irradiated 3T3 fibroblasts treated with CQ 7 days after RT. V=vacuole-like structure, M=mitochondria, LD=lipid droplet, ER=endoplasmic reticulum. Black arrows indicate higher magnification of mitochondria. Scale bar is 1 µm. **(B)** Representative IF images of ATP5A1 (green) and nuclear (blue) staining 7 days after RT with and without CQ treatment in irradiated and unirradiated 3T3 fibroblasts. Scale bars are 10 µm. **(C)** Quantification of the mitochondrial aspect ratio at 3 and 7 days after RT in unirradiated and irradiated 3T3 fibroblasts with and without CQ. Each data point represents mitochondria from one individual cell (n=3). **(D)** Representative Western blot of ATG5 protein expression compared to GAPDH loading control in irradiated and unirradiated shScr and shAtg5 3T3 fibroblasts. **(E)** Representative Western blot of LC3B-I and LC3B-II protein expression compared to α-tubulin loading control in irradiated and unirradiated shScr and shAtg5 fibroblasts 7 days after RT. **(F)** Relative gene expression of *Mfn1, Mfn2, Opa1,* and *Sirt5* in shScr and shAtg5 fibroblasts 3 and 7 days after RT (n=3). **(G)** Representative IF images of ATP5A1 (green) and nuclear (blue) staining at 7 days after RT in irradiated and unirradiated shScr and shAtg5 fibroblasts. Scale bars are 10 µm. **(H)** Quantification of the mitochondrial aspect ratio at 3 and 7 days after RT in unirradiated and irradiated shScr and shAtg5 fibroblasts. Each data point represents mitochondria from one individual cell (n=3). For **C** and **H**, statistical analysis was determined by one-way ANOVA with Games-Howell pairwise comparison corrections for non-equal variances with *p<0.05, **p<0.01, and ***p<0.001. Error bars show standard deviation.

Next, we analyzed mitochondrial fusion-related genes at 3 and 7 days after RT in shAtg5 cells. In our wildtype cells, we saw a moderate increase in mitofusin gene expression at early timepoints, and a significant increase in *Sirt5* at 7 days after RT (**Fig. 1F**). We saw similar trends in shScr cells, but these increases were largely abrogated in shAtg5 cells (**Fig. 3F**). Finally, we analyzed the mitochondrial aspect ratio from ATP5A1 IF staining (**Fig. 3G**) and confirmed that elongated mitochondrial morphology was reduced in irradiated shAtg5 fibroblasts (**Fig. 3H**). We also probed mitophagy through Western blotting of phosphorylated PTEN-induced kinase 1 (PINK1) in irradiated and unirradiated cells with CQ incubation^66^. Equivalent levels of phospho-PINK1 between irradiated and unirradiated cells were observed as well as a decreasing trend of expression for both conditions following CQ incubation, suggesting that mitochondria are not being targeted for degradation (**Fig. S1F–G**). These results indicate that radiation induces increased autophagic flux in fibroblasts, causing mitochondria to undergo fusion and elongate to prevent being degraded through bulk autophagy.

### Metabolic flux analysis reveals that increased mitochondrial respiration is linked to autophagy and FAO

To further explore mitochondrial changes, we stained fibroblasts with MitoTracker Deep Red and found an increase in intensity, especially 7 days after RT (**Fig. 4A**, **S2A**). Increased staining at the whole cell level could be the result of either increased mitochondrial mass, increased membrane potential, or a combination^67,68^. Short-term autophagy inhibition through CQ dramatically increased MitoTracker Deep Red intensity for both irradiated and unirradiated cells (**Fig. 4A**). While we saw a strong increase in MitoTracker Deep Red intensity for irradiated 7-day post-RT shScr cells, there was no further increase in MitoTracker Deep Red intensity after radiation when autophagy was blocked long-term in shAtg5 cells (**Fig. 4B**). To validate our findings, we performed staining with tetramethylrhodamine ethyl ester (TMRE) to evaluate membrane potential at the individual mitochondria level^69^. No significant changes in TMRE intensity were observed between untreated control and irradiated cells at either early or late time points after RT (**Fig. 4C**, **S2B**), suggesting increased MitoTracker Deep Red staining is due to increased total area of mitochondria per cell as opposed to increased localized membrane potential. In CQ-treated cells, both irradiated and unirradiated fibroblasts demonstrated a stark increase in membrane potential, which was not observed in shAtg5 knockdown cells (**Fig. 4D**, **S2C**). Although these two methods for blocking autophagy show similarities in preventing mitochondrial elongation, these results suggest that they may have different impacts on mitochondrial bioenergetics and respiration.

**Figure 4.**
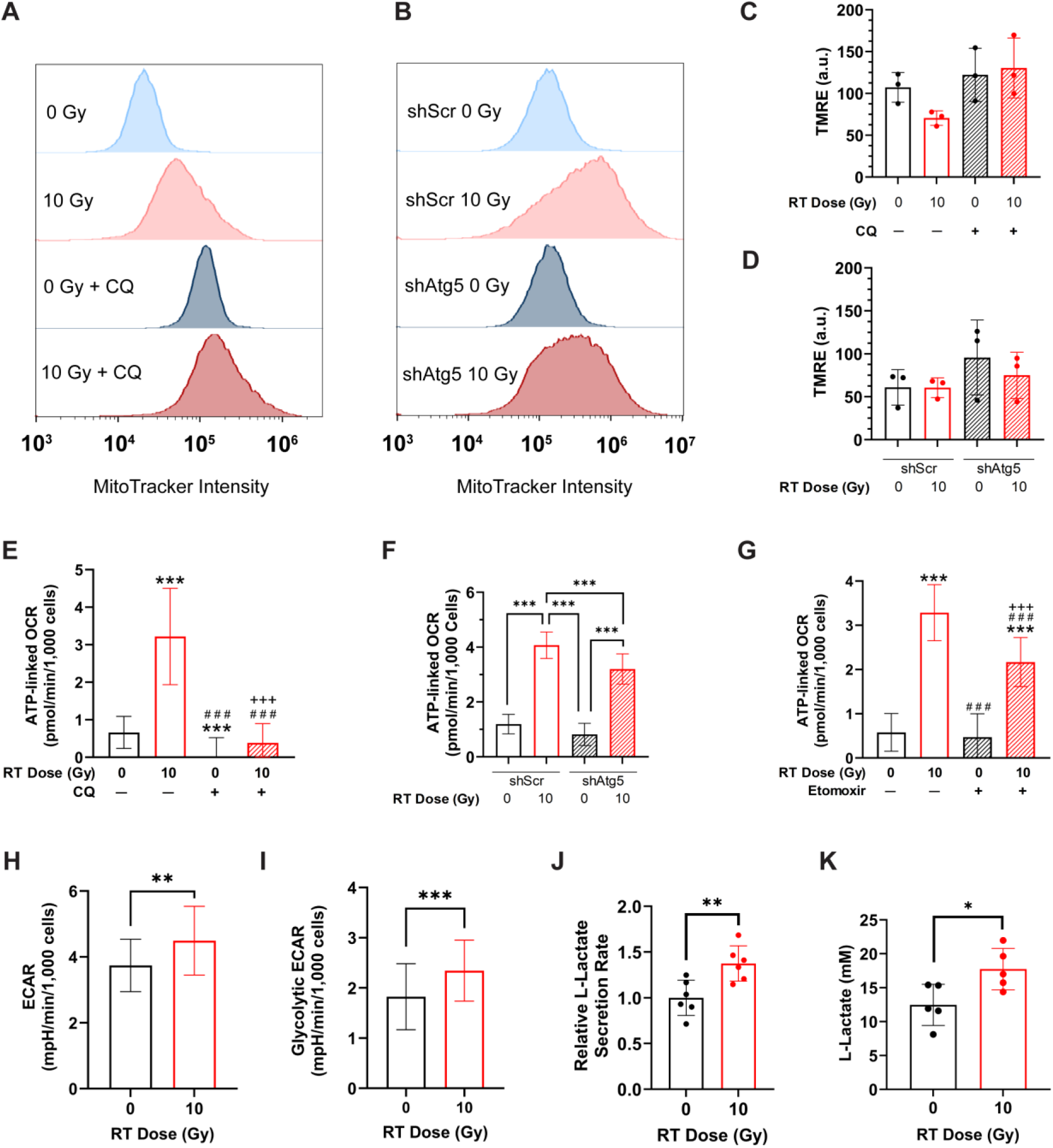
Radiation-induced autophagy maintains high levels of mitochondrial respiration. Representative flow cytometry analysis of irradiated and unirradiated 3T3 fibroblasts stained with MitoTracker Deep Red **(A)** with and without CQ incubation and **(B)** in shScr and shAtg5 fibroblasts 7 days after RT. Tetramethylrhodamine ethyl ester (TMRE) intensity measurements of 3T3 fibroblasts **(C)** with and without CQ incubation (n=3) and **(D)** in shScr and shAtg5 fibroblasts 7 days after RT (n=3). ATP-linked oxygen consumption rate (OCR) in irradiated and unirradiated 3T3s **(E)** with and without CQ incubation (n=3; non-equal variances) and **(F)** in shScr and shAtg5 fibroblasts 7 days after RT (n=3; equal variances). **(G)** ATP-linked OCR in irradiated and unirradiated 3T3s 7 days after RT with and without etomoxir incubation to evaluate fatty acid oxidation (FAO) (n=3; equal variances). **(H)** Baseline extracellular acidification rate (ECAR) in irradiated and unirradiated 3T3s 7 days after RT (n=3). **(I)** Glycolytic ECAR in irradiated and unirradiated 3T3s 7 days after RT (n=3). **(J)** Relative L-lactate secretion rate in irradiated and unirradiated 7-day post-RT 3T3 fibroblasts (n=6). **(K)** Quantification of L-lactate concentration in fibroblast CM collected 7 days after RT from irradiated and unirradiated 3T3s (n=4). Statistical analyses for **E – G** determined by one-way ANOVA using either Tukey or Games-Howell pairwise comparison corrections for equal or non-equal variances, respectively. Symbols for pairwise comparisons: * for comparison to 0 Gy & 0 µM inhibitor, # for comparison to 10 Gy & 0 µM inhibitor, and + for comparison to 0 Gy & high concentration of inhibitor, where *, ^#^, ^+^p<0.05, **, ^##^, ^++^p<0.01, and ***, ^###^, ^+++^p<0.001. For **H – K**, statistical analyses determined by unpaired two-tailed t-tests. All error bars show standard deviation.

We next analyzed mitochondrial respiration through a mitochondrial stress test (**Fig. S2D– E**). Irradiated fibroblasts had a roughly 2.5-fold increase in ATP-linked oxygen consumption rate (OCR) compared to unirradiated fibroblasts 3 days after RT (**Fig. S2F**) and greater than a 4.5-fold increase at the 7-day timepoint (**Fig. 4E**), matching observations of increased MitoTracker staining despite minimal differences in membrane potential between irradiated and unirradiated cells. Autophagy inhibition by CQ completely abrogated the large ATP-linked OCR increase in irradiated fibroblasts. When analyzing mitochondrial respiration in irradiated shAtg5 cells, we saw a significant decrease in ATP-linked OCR – approximately 25% – compared to shScr cells (**Fig. 4F**). These results demonstrate that irradiated fibroblasts are highly aerobic, but blocking autophagy through CQ treatment has a more severe effect on mitochondrial respiration than genetic knockdown.

Since our gene expression analysis demonstrated increased expression of *Sirt5* in irradiated fibroblasts (**Fig. 1F**) and our TEM results additionally indicated high levels of lipid droplets in irradiated cells (**Fig. 1A**), we hypothesized that altered lipid metabolism influences radiation-induced mitochondrial respiration. We confirmed increased neutral lipid accumulation in irradiated fibroblasts through both perilipin-2 IF (**Fig. S3A–B**) and Oil Red O (ORO, **Fig. S3C– D**) staining. We also observed increased fatty acid uptake as determined by fluorescence imaging of cells after incubation with BODIPY-conjugated palmitate (**Fig. S3E–F**). These results suggest that FAO supports mitochondrial respiration in irradiated fibroblasts even with increased neutral lipid stores^70,71^. We confirmed this by performing a mitochondrial stress test with irradiated and unirradiated fibroblasts following incubation with etomoxir (**Fig S2D–E**), where we approximately saw a 40% decrease in ATP-linked OCR in irradiated cells when FAO is blocked at 3- and 7-days post-RT (**Fig. 4G**, **S3G**). Upon autophagy inhibition with CQ, we observed a stark decrease in lipid droplets in irradiated fibroblasts that was confirmed through both perilipin-2 (**Fig. S3H**) and ORO (**Fig. S3I**) staining. Similar trends in neutral lipid storage were also observed in human iMFs (**Fig. S3J**). These results demonstrate that irradiated fibroblasts have an altered lipid metabolic profile that could partially be regulated by autophagy.

We also observed an increase in baseline extracellular acidification rate (ECAR) only in irradiated fibroblasts after 7 days, suggesting an increase in aerobic glycolysis despite high mitochondrial respiration (**Fig. 4H**, **S4A**). We confirmed that this increased ECAR was due to glycolysis by removing pyruvate, used in the mitochondrial stress when evaluating baseline ECAR which can confound glycolytic ECAR results, and monitoring the change in ECAR following glucose injection (**Fig. 4I**, **S4B–C**). To further validate increased glycolysis in irradiated cells, we confirmed an increase in lactate secretion rate of irradiated cells compared to unirradiated controls (**Fig. 4J**) and found a 5 mM increase in the lactate concentration in the conditioned media (CM) of 7-day post-RT fibroblasts compared to unirradiated controls (**Fig. 4K**). Together, we show that irradiated fibroblasts have an altered metabolic profile that is partially regulated by increased autophagic flux.

### Irradiated fibroblasts demonstrate a mitochondrial stress response that is disrupted following autophagy inhibition

Our respiration data indicated an increase in non-mitochondrial OCR that increased with time after RT (**Fig. 5A**). These results suggest increased cellular and mitochondrial stress and increased ROS generation^72–74^. Through MitoSOx flow cytometry analysis, we saw an increase in the presence of mitochondrial superoxide generation in irradiated cells but only after 7 days (**Fig. 5B**). We also found a strong shift in mitochondrial superoxide production in both irradiated and unirradiated cells upon autophagy inhibition with CQ but did not observe this shift upon autophagy inhibition via genetic knockdown (**Fig. 5C**), supporting our observation that CQ incubation causes a larger decrease in mitochondrial respiration compared to genetic knockdown of ATG5. These data suggest a mitochondrial stress response in irradiated fibroblasts that increases over time.

**Figure 5.**
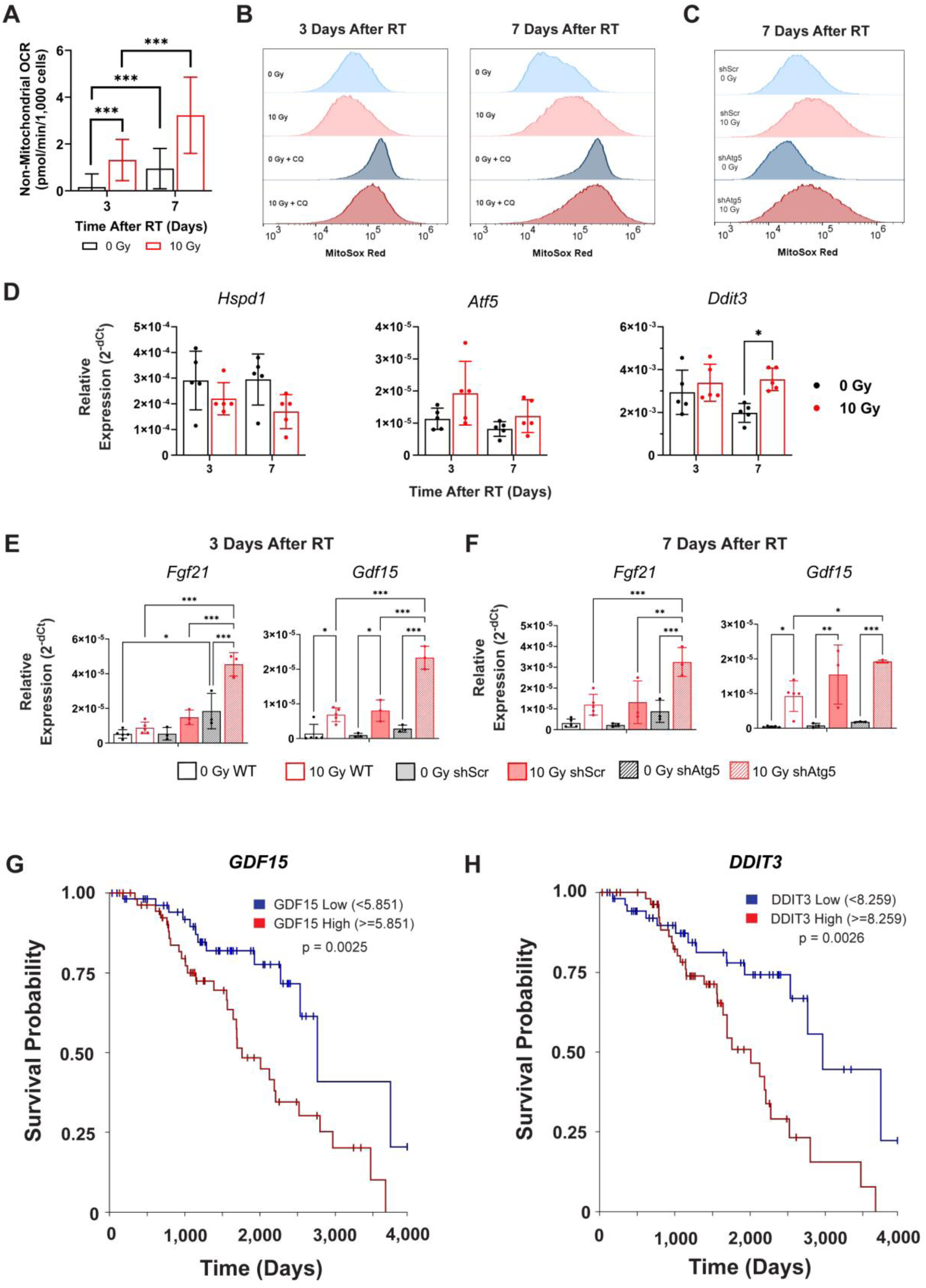
Irradiated fibroblasts demonstrate hallmarks of a mitochondrial stress response that is regulated by autophagy. **(A)** Non-mitochondrial OCR in irradiated and unirradiated 3T3s at 3 and 7 days after RT (n=3). **(B)** Representative flow cytometry histogram analysis of MitoSOX staining in unirradiated and irradiated 3T3 fibroblasts 3 and 7 days after RT. **(C)** Representative flow cytometry histogram analysis of MitoSOX staining in unirradiated and irradiated shScr and shAtg5 cells 7 days after RT. **(D)** Relative gene expression of *Hspd1, Atf5*, and *Ddit3* in control and irradiated 3T3 fibroblasts at 3 and 7 days after RT (n=5). Relative gene expression of *Fgf21* and *Gdf15* at **(E)** 3 and **(F)** 7 days after RT in wild type, shScr, and shAtg5 fibroblasts (n=3–5). Kaplan-Meier analysis of overall survival utilizing normal adjacent tissue (NAT) data from the TCGA BRCA dataset accessed through UCSC’s Xena platform^81,82^ and grouping patients based on high or low gene expression of **(G)** *GDF15* and **(H)** *DDIT3*. For **A**, statistical analysis was performed using a one-way ANOVA with Tukey pairwise comparison corrections, with ***p<0.001. For **D**, statistical analysis was determined by a 2-way ANOVA with multiple comparisons by the Sidak method with *p<0.05. For **E** and **F**, a one-way ANOVA with multiple comparisons by the Sidak method was performed, with*p<0.05, **p<0.01, and ***p<0.001. For **G** and **H**, statistical analysis was performed using the log-rank test. Error bars show standard deviation.

We then evaluated the expression of genes involved in the mitochondrial stress response^75,76^. We saw a trend of decreasing *Hspd1* gene expression and increasing expression of *Atf5* and *Ddit3* (**Fig. 5D**). We also observed a strong upregulation of *Fgf21* and *Gdf15*, which are critical mitokines observed under mitochondrial stress^77,78^ (**Fig. 5E–F**). We observed similar trends in mitokine expression in human iMFs at 7 days after RT (**Fig. S5A**). When we analyzed mitokine gene expression under autophagy inhibition through ATG5 knockdown, expression was further increased at both early and late timepoints following radiation (**Fig. 5E–F**). These mitokines have been shown to influence tumor cell behavior^79,80^ but are also typically upregulated in conjunction with other cytokines and chemokines. We therefore additionally performed a multiplex cytokine assay on CM from irradiated fibroblasts collected after 7 days (**Fig. S5B, Table S1**). Multiple cytokines and chemokines were increased in fibroblast CM following radiation, with interleukin-6, chemokine (C-X-C motif) ligand 1 (CXCL1), vascular endothelial growth factor (VEGF), and C-C motif ligand 2 (CCL2) being the top four hits. Gene expression analysis of these factors after RT as well as after autophagy inhibition through ATG5 knockdown showed significantly decreased *Il6* yet no significant changes in *Ccl2*, *Vegfa*, *or Cxcl1* in the irradiated shAtg5 cells compared to scramble controls, although there was an increasing trend in *Ccl2* expression (**Fig. S5C**). Taken together, these results support a mitochondrial stress response in fibroblasts following RT that is disrupted when autophagy is blocked, resulting in complex changes in expression of cytokines, chemokines, and mitokines that may influence TNBC cells in the context of recurrence following RT.

### Irradiated fibroblasts induce an aggressive TNBC cell phenotype

Through The Cancer Genome Atlas Breast Invasive Carcinoma (TCGA BRCA) dataset^81,82^, we evaluated normal adjacent tissue (NAT) collected and analyzed at the time of surgery. Kaplan-Meier analysis of overall survival with patients grouped based on high or low expression of *GDF15* and *DDIT3* showed that patients with high expression of either gene in the NAT had significantly lower overall survival (**Fig. 5G–H**), which highlights the potential of the mitochondrial stress response in fibroblasts influencing TNBC recurrence mechanisms.

To explore the impact of irradiated fibroblasts on TNBC cell behavior, we utilized a scratch assay where TNBC cells were exposed to either irradiated or unirradiated fibroblast CM, and the gap closure was observed over a 15-hour incubation period (**Fig. 6A–B**). TNBC cells incubated in irradiated fibroblast CM closed a greater percentage of the gap across the 15-hour time period (**Fig. 6C**). These outcomes were replicated in a TNBC-fibroblast co-culture assay (**Fig. 6D**), where TNBC cells co-cultured with irradiated fibroblasts closed a greater percentage of the gap compared to those cultured with unirradiated fibroblasts (**Fig. 6E–F)**. These results demonstrate a link between irradiated fibroblasts and the induction of an aggressive migratory phenotype in TNBC cells.

**Figure 6.**
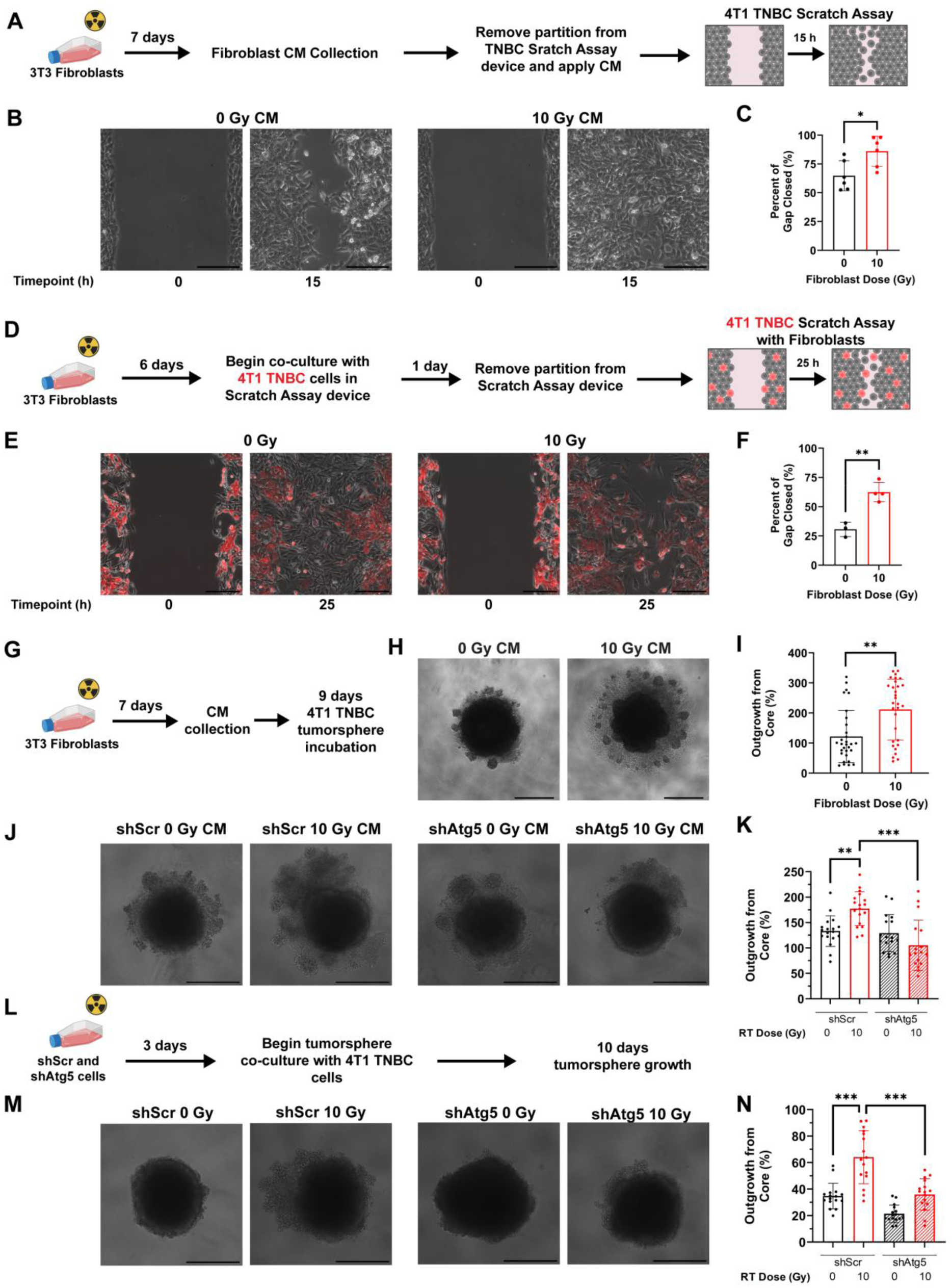
Irradiated fibroblasts induce an aggressive phenotype in TNBC cells that is disrupted upon fibroblast autophagy inhibition. (**A**) Schematic representing time course of conditioned media (CM) collection for a 4T1 TNBC cell scratch wound assay. (**B**) Representative images of 4T1 TNBC cells in scratch wound assays at 0- and 15-hour timepoints following incubation in either irradiated or unirradiated control fibroblast CM. The corresponding quantification (n=6) is shown in (**C**). (**D**) Schematic representing time course of scratch wound assay with 4T1 TNBC cells co-cultured with either irradiated or unirradiated 3T3 fibroblasts. (**E**) Representative images of 4T1 TNBC cells (red) in scratch wound assays co-cultured with irradiated or unirradiated 3T3 fibroblasts (unlabeled) at 0- and 25-hour timepoints. The corresponding quantification (n=3–4) is shown in (**F**). (**G**) Schematic representing time course of CM collection for 3D 4T1 TNBC cell tumorsphere assay. (**H**) Representative images of 4T1 TNBC tumorspheres after 9 days of incubation in either irradiated or unirradiated control fibroblast CM. The corresponding quantification (n=30 tumorspheres) is shown in (**I**). (**J**) Representative images of 4T1 TNBC tumor spheroids after 9 days of incubation in either irradiated or unirradiated shScr or shAtg5 fibroblast CM. The corresponding quantification (n=18 tumorspheres) is shown in (**K**). (**L**) Schematic representing time course of shScr and shAtg5 fibroblast irradiation and co-culture with 4T1 TNBC cells in tumorsphere assay. (**M**) Representative images of 4T1 TNBC tumorspheres after 10 days of co-culture with irradiated or unirradiated shScr or shAtg5 fibroblasts. The corresponding quantification (n=16 tumorspheres) is shown in (**N**). Statistical analysis was performed using the following: for **C** and **F**, an unpaired two-tailed t-test; for **I**, a Mann-Whitney test for non-normally distributed distributions; for **K** and **N**, a one-way ANOVA with Tukey and Games-Howell pairwise comparison corrections, respectively. *p<0.05, **p<0.01, and ***p<0.001. Scale bars are 200 µm in **B** and **E** and 500 µm in **H**, **J**, and **M**. Error bars show standard deviation.

### Blocking autophagy and disrupting the mitochondrial stress response in irradiated fibroblasts inhibits TNBC tumorsphere growth

We next evaluated this pro-metastatic behavior in a more biologically relevant 3D system. Since CTCs have been shown to survive detached from the extracellular matrix due to increased mitochondrial respiration^33,34^, we evaluated how irradiated fibroblasts alter tumor spheroid growth. We observed a stark increase in TNBC tumorsphere outgrowth from the dense center when incubated with CM from irradiated fibroblasts after 9 days (**Fig. 6G–I**), mirroring CTC survival behavior. Since we previously saw that blocking autophagy in irradiated fibroblasts impacts mitochondrial dynamics and the mitochondrial stress response, we again utilized the tumorsphere analysis with shScr and shAtg5 fibroblast CM. Incubation with shScr CM from irradiated cells also resulted in a significant outgrowth in TNBC spheroids, but outgrowth decreased significantly with CM from irradiated shAtg5 cells (**Fig. 6J–K**). These results were validated using a co-culture tumorsphere system where at 3-days post-RT, irradiated and unirradiated shScr and shAtg5 fibroblasts were incubated with TNBC cells (**Fig. 6L–N**).

These studies identify an important relationship between metabolic reprogramming in fibroblasts, autophagy, and a mitochondrial stress response (**Fig. 7**). We demonstrate that blocking autophagic flux in irradiated fibroblasts disrupts the mitochondrial stress response and may be beneficial in reducing aggressive and metastatic TNBC behavior in the post-irradiated microenvironment.

**Figure 7.**
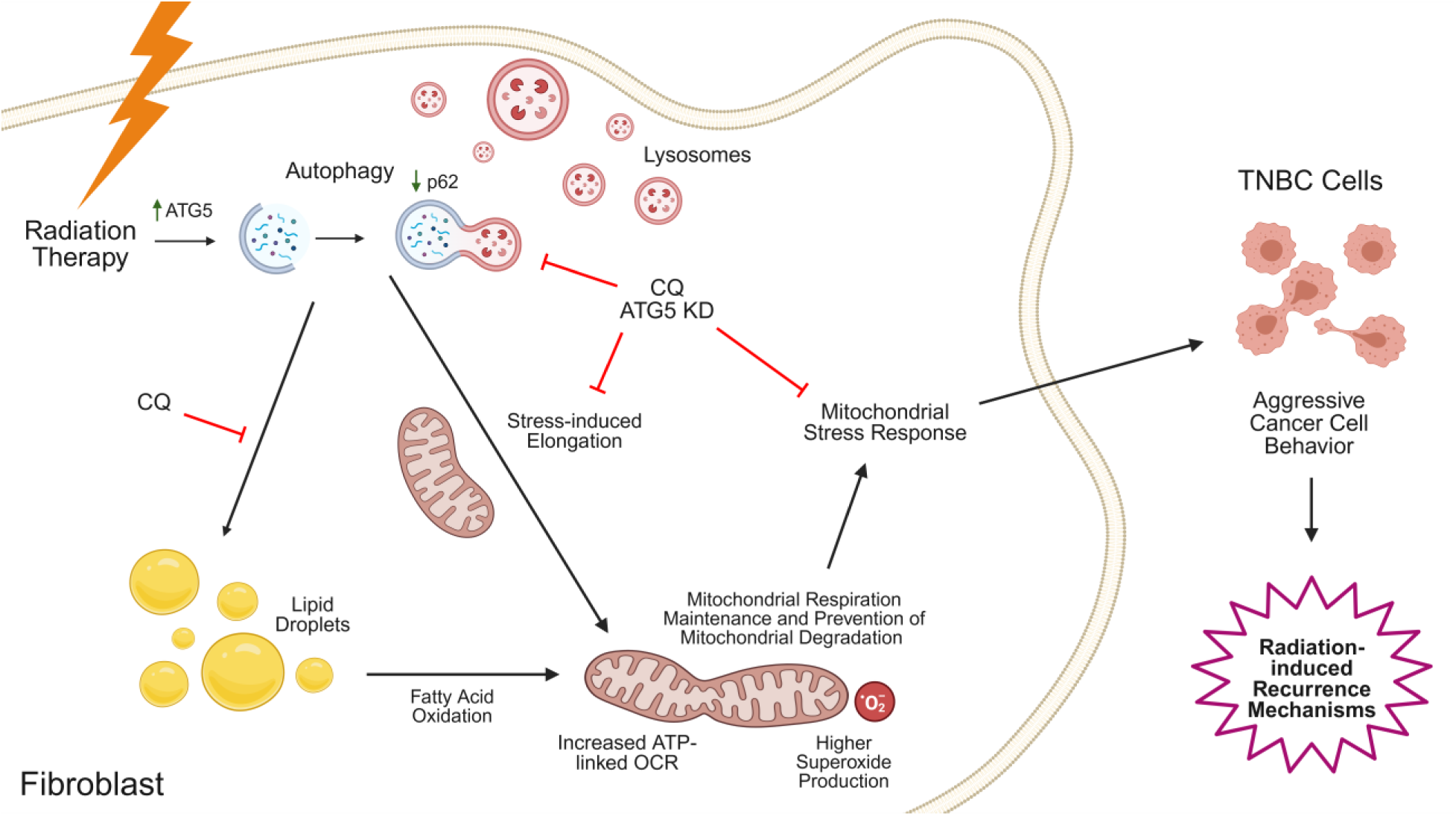

## Discussion

In this study, we establish that irradiated fibroblasts undergo extensive metabolic reprogramming up to 7 days following RT that involves mitochondrial morphology elongation, upregulation of autophagy, and increased mitochondrial respiration, aerobic glycolysis, and FAO. We uncover a resulting mitochondrial stress response that is disrupted upon autophagy inhibition. We demonstrate that irradiated fibroblasts can induce an aggressive phenotype in TNBC cells as part of this response through both secreted factors and direct physical interaction. By blocking autophagy in irradiated fibroblasts, we observe an abrogation of increased outgrowth in detached TNBC tumorspheres, suggesting a previously unappreciated link between the mitochondrial stress response and autophagic flux in irradiated fibroblasts and TNBC recurrence.

### Prolonged autophagy upregulation following RT – friend or foe?

Radiation-induced autophagy can be a survival mechanism but can also induce cell death.^83^ In addition to these two disparate pathways, autophagy can also become defective in irradiated cells, suggesting that the exact role autophagy plays following RT may be cell type and context dependent^84^. It has been demonstrated that increased autophagy following radiation at lower total doses and on the time scale up to 24 hours post-RT can provide a cell survival and protective effect^85^. Multiple downstream effects of radiation could activate autophagy, with many studies suggesting that this is due to increased endoplasmic reticulum stress and the activation of the unfolded protein response^86,87^. Our results of increased levels of autophagic flux at both 3- and 7-days following radiation are surprising as persistent upregulation of autophagic flux and prolonged autophagy has been shown to promote cell death^88^. An extended level of high autophagic flux in irradiated fibroblasts may be important in TNBC recurrence, where the recruitment of CTCs to the irradiated microenvironment has been shown to play a role in pre-clinical recurrence mechanisms^16,17^. These outcomes are significantly impacted by immune cells, especially macrophages^89,90^. Autophagy has been observed in unirradiated cells through radiation-induced bystander effects^91–93^, and increased autophagy in macrophages may promote a pro-tumor phenotypic shift as macrophages with decreased levels of autophagy have been shown to exhibit a pro-inflammatory and anti-tumor phenotype^94,95^. These studies support our findings that increased autophagic flux in irradiated fibroblasts and the resultant crosstalk with other cells in the breast microenvironment may be important mechanisms in TNBC recurrence.

### Mitochondrial elongation as a protective mechanism to maintain respiration under high levels of autophagy

High doses of radiation have been shown to increase mitochondrial superoxide production after 24 hours, which has been attributed to a reduction in glutathione peroxidase activity^85^. Increased superoxide production typically results in increased mitochondrial damage and PINK-1 dependent mitophagy,^96^ and increased mitophagy has been reported following radiation^97^. We did not see an increase in phospho-PINK1 expression at 3 or 7 days after RT, indicating a lack of mitophagy, and additionally measured high levels of mitochondrial superoxide production after 7 days. The high levels of non-mitochondrial OCR observed in irradiated fibroblasts may suggest high levels of intracellular oxidases which could further promote oxidative damage to organelles including mitochondria^72–74^. However, the RT response undoubtedly requires high levels of ATP production to allow for cell survival and will require high levels of mitochondrial respiration^98–100^, supporting our findings of increased ATP-linked OCR following radiation. Under starvation-induced autophagy, mitochondrial elongation and increased fusion protein expression has been shown to protect critical mitochondria that are needed to sustain respiration and ATP production from degradation^101^. Given our findings of increased mitochondrial elongation up to 7 days after RT coupled with increased mitochondrial respiration, our results suggest that protecting mitochondria from degradation by autophagy may be necessary to maintain high ATP levels for the damage response and allow for cell survival despite their role in increased cellular stress and oxidative damage.

### The mitochondrial stress response following radiation is an important and underappreciated post-therapy mechanism that could influence TNBC recurrence

Mitochondrial stress responses have recently been recognized as important quality control processes that are critical in cancer progression^29,102,103^. Two of these processes, the mitochondrial unfolded protein response and the mitochondrial integrated stress response, are closely linked and the downstream effects of their activation can exhibit both an autocrine and paracrine effect on the microenvironment^26–28^. Two mitokines that are commonly upregulated during these stress responses are GDF15 and FGF21, and they have been shown as biomarkers following radiation^104–106^ as well as other disease states like obesity and insulin resistance that are prone to mitochondrial damage and stress^107^. The secretion of these mitokines have significant metabolic effects on recipient cells^108^, which agrees with our TCGA analysis from NAT. How blocking autophagy influences the mitochondrial stress response is understudied, especially after radiation, and we demonstrate that blocking autophagy leads to an increased mitokine expression profile that mitigates aggressive tumor cell behavior^77^. In addition, lactate secretion has long been identified as an important metabolite in the tumor microenvironment^109^, and lactate metabolism is closely related to mitochondrial stress and dysfunction^110,111^. Acidosis and increased lactate secretion in the tumor microenvironment is known to influence immune and tumor cell behavior, and CAFs secrete lactate once activated^112–115^. We demonstrate that irradiated fibroblasts increase lactate secretion in addition to increased mitochondrial respiration, highlighting the importance of the secretory profile of irradiated fibroblasts in post-RT recurrence and treatment failure.

### Blocking autophagy as a potential combination therapy with ionizing RT

As of early 2025, there are four active clinical trials exploring the use of hydroxychloroquine, a derivative of CQ, for the treatment of cancer patients, with one trial including breast cancer patients^116^. In these and past clinical trials, hydroxychloroquine is largely administered with chemotherapeutics and antimetabolites although studies have analyzed combinatorial effects with hormone therapies and monoclonal antibodies to target treatment-resistant and dormant tumor cells^117,118^. There has been little attention to the use of CQ or its derivatives in combination with RT. An early trial utilized CQ successfully in glioblastoma multiforme patients following surgical intervention and in conjunction with conventional chemotherapy and RT^119^. Pre-clinically, CQ has been used in combination with RT for breast tumors in mice to engage the immune system to target the primary tumor^120^. The role of treated normal tissue cells and the potential mechanisms of how stromal cells like fibroblasts could influence survival and recurrence rates following RT and in conjunction with CQ have not been evaluated. These studies in combination with our data suggest the potential viability of CQ usage to promote autophagy inhibition as a combinatorial therapy following RT in TNBC.

In summary, we demonstrate an essential relationship between mitochondrial dynamics, autophagy, and a mitochondrial stress response in fibroblasts that alters TNBC cell behavior following radiation. Our study provides a link between metabolic profiles in fibroblasts and an aggressive TNBC phenotype that may lead to treatment failure following RT and impact TNBC patient survival. Our work suggests that targeting the crosstalk between irradiated fibroblasts and TNBC cells may reduce the risk of recurrent disease and improve outcomes for TNBC patients who undergo RT.

### Limitations of the study

We blocked autophagy and showed how this exacerbates the mitochondrial stress response in fibroblasts, and future studies are needed to determine the exact mechanisms that drive aggressive TNBC behavior following RT. In addition, CTCs may have an altered metabolic profile compared to primary tumor cells and may exhibit unique responses to irradiated fibroblasts. Although we show blocking autophagy in irradiated fibroblasts abrogates an aggressive TNBC phenotype, additional studies should probe how the introduction of TNBC cells to irradiated fibroblasts modifies the mitochondrial stress response. The incorporation of other cell types typically found in the post-RT microenvironment in breast cancer should also be evaluated. For example, adipocytes, another abundant cell type in the breast tissue microenvironment, have been demonstrated to influence the fibroblast radiation response^121^. Adipocyte lipid metabolic response to radiation may alter extracellular fatty acid availability for fibroblasts, impacting autophagy and lipid metabolism in irradiated fibroblasts. This could undoubtedly shape the mitochondrial stress response in fibroblasts. Evaluating these and other cell types will be critical for elucidating mechanisms of TNBC recurrence.

## Acknowledgments

The authors thank Dr. Christopher Contag for providing luciferase-labeled 4T1s, Dr. Charlotte Kuperwasser for providing immortalized reduction mammoplasty fibroblast cells, Dr. Craig Duvall for the use of the IVIS, and Dr. Michael Freeman for the use of the Cesium irradiator. The results shown here are in part based upon data generated by the TCGA Research Network: https://www.cancer.gov/tcga. We would also like to thank Dr. Rachelle Johnson for her guidance in accessing and analyzing TCGA BRCA data, and Dr. Alyssa Hasty for her critical review of our manuscript. We would also like to thank the Translational Pathology Shared Resource core facility for ex vivo sample preparation. Experiments using the Seahorse XFe96 Metabolic Flux Analyzer were performed in the Vanderbilt High-Throughput Screening Core Facility, which receives support from the Vanderbilt Institute of Chemical Biology and the Vanderbilt Ingram Cancer Center (P30 CA68485). TEM was performed in part through the use of the Vanderbilt Cell Imaging Shared Resource, supported by NIH grants CA68485, DK20593, DK58404, DK59637, EY08126, and 1S10OD034315-01. This research was financially supported by National Institutes of Health grant R00CA201304 (MR), the American Cancer Society Research Scholar Grant RSG-21-151-01-CDP (MR), the Ruth L. Kirschstein National Research Service Award grant T32DK101003 (KCC), and the Maximizing Access to Research Careers Award grant T34GM136451 (LSB). Schematics were prepared using BioRender.

## Author Contributions

Conceptualization: KCC, MR

Methodology: KCC, SEM, VKM, JDY, VG, MR

Investigation: KCC, SEM, VKM, LSB, KMS YKM, YII, AAG, SAW, TZ, EK

Visualization: KCC

Supervision: MR

Writing—original draft: KCC

Writing—review & editing: KCC, SEM, VG, MR

## STAR Methods

### Cell culture

NIH 3T3 murine embryonic fibroblasts were obtained from ATCC (CRL-1658), and immortalized human reduction mammoplasty fibroblasts (iMFs) were obtained from Dr. Charlotte Kuperwasser (Tufts University) in April 2022. Fibroblasts were cultured in high glucose DMEM (Gibco 11995-065) containing 10% bovine calf serum and 1% penicillin/streptomycin. 4T1 TNBC cells were obtained from ATCC and cultured in RPMI 1640 medium (Gibco 11875-093) containing 10% heat-inactivated fetal bovine serum and 1% penicillin/streptomycin. All cells were maintained at 37°C in a humified incubator containing 5% CO_2_ and were routinely evaluated for the presence of mycoplasma using the Myco-Sniff-Rapid™ Mycoplasma Luciferase Detection Kit (MP Biomedicals 0930504-CF).

### ATG5 knockdown cell line generation and maintenance

The shRNA constructs of ATG5 and scramble control were obtained from Sigma Aldrich in either a pLKO.1 or pLKO.2 backbone. Target sequences and plasmid backbone for each construct were as follows: shScramble (SHC002, pLKO.1): 5’-CCGGCAACAAGATGAAGAGCACCAACTCGAGTTGGTGCTCTTCATCTTGTTGTTTTT-3’ shATG5 (TRCN0000375819, pLKO.2): 5’-AGCCGAAGCCTTTGCTCAATG-3’

Each of the plasmids were amplified in DH5α E. coli and purified using QIAprep Spin Miniprep Kit (Qiagen 27104). Following amplification, 1 µg of each plasmid was transfected into HEK293T cells using FuGene6 (Promega E2691) along with 0.75 µg packaging plasmid (psPAX2; AddGene 12260) and 0.25 µg envelope plasmid (pMD2.G; AddGene 12259). Media containing lentiviral particles (LVP) was harvested after 48 hours and filtered through a 0.45 µm syringe filter (Fisher Scientific 09-928-061) before being aliquoted. LVP aliquots were either flash frozen in LN_2_ or immediately transduced into target cells. For LVP transduction, viral dilutions ranging from 1:5 to 1:500 in 500 µL of polybrene-containing, penicillin/streptomycin-free complete media was added to a 6-well plate, followed by 50,000 3T3 cells. LVPs containing the shRNA construct of interest were diluted in polybrene-containing, penicillin/streptomycin-free complete media. After 48 hours, the media was exchanged for puromycin-containing complete media. Puromycin concentration of 2 µg/mL was used for selection in 3T3 cells. Puromycin selection of polyclonal cells continued for 1 week or until cells were confluent. Cells were maintained in basal media with 0.5 µg/mL puromycin until radiation experiments were performed, at which point puromycin was removed from the media. A polyclonal population was used for knockdown of ATG5 as recommended to maintain reduction of autophagic flux^122^. Knockdown efficiency and reduction of autophagic flux was confirmed by Western blotting for ATG5 and LC3B.

### Animals

Animal studies were performed in accordance with institutional guidelines and protocols approved by the Vanderbilt University Institutional Animal Care and Use Committee. 8-10-week-old female BALB/c mice were obtained from Charles River Laboratories. Mice were allowed free access to standard diet and water and maintained under a 12 h light/12 h dark cycle.

### Radiation

For cell experiments, cells were grown to 70% confluency in T300, T75, or T25 flasks and then irradiated to a dose of 10 Gy using a cesium source. Following irradiation, cells were immediately passaged into new plates or flasks with fresh media. Cell counts at the indicated post-RT timepoints were kept constant between irradiated and non-irradiated cells. For mammary fat pad (MFP) radiation experiments, MFPs were harvested from female BALB/c mice after sacrifice with CO_2_ asphyxiation followed by cervical dislocation. Tissues were then irradiated to a dose of 20 Gy *ex vivo* using a cesium source. MFPs were cultured in complete DMEM media at 37°C and 5% CO_2_ for seven days before fixing in neutral buffered formalin (VWR 16004-128) overnight at 4°C. Tissues were then triple rinsed in PBS with gentle rocking and placed in 70% ethanol in preparation for processing. Samples were embedded into paraffin blocks and sectioned (5 µm) onto glass microscope slides by the Vanderbilt Translational Pathology Shared Resource core facility.

### Transmission electron microscopy

Experiments were performed by the Vanderbilt Cell Imaging Shared Resource core facility. All electron microscopy reagents were purchased from Electron Microscopy Sciences unless noted otherwise. At 7 days after RT, 3T3 fibroblasts with and without 100 µM CQ treatment for 6 hours were fixed at 70% confluency with warmed 2.5% glutaraldehyde in 0.1 M cacodylate buffer for 1 hour at room temperature followed by 24 hours at 4°C. After fixation, the cells were mechanically lifted from the tissue culture plates and pelleted, then sequentially postfixed with 1% tannic acid, 1% osmium tetroxide, and stained en bloc with 1% uranyl acetate. Samples were then dehydrated in a graded ethanol series, infiltrated with Quetol 651–based Spurr’s resin, and polymerized at 60°C for 48 hours. Ultrathin sections were prepared on a UC7 ultramicrotome (Leica) with a nominal thickness of 70 nm and collected onto 300-mesh nickel grids. Sections were stained with 2% uranyl acetate and lead citrate and imaged using a Tecnai T12 operating at 100 kV equipped with an AMT NanoSprint CMOS camera using AMT imaging software.

### Fibroblast CM and lactate concentration

Irradiated and unirradiated 3T3s were plated in T25 flasks in complete growth media for 7 days, at which point supernatant was collected and filtered through a 0.2 μm filter to remove cell debris. Cell counts between irradiated and unirradiated flasks at the time of CM collection were kept constant. Filtered CM was stored at -80°C until use. L-lactate concentration in CM was determined using Cayman Chemical’s L-Lactate Assay Kit (700510) following the manufacturer’s instructions. When 4T1s were incubated in fibroblast CM, complete media was removed 24 hours prior to 70% confluency, washed with phosphate buffered saline (PBS), and replaced with CM for incubation at 37°C. For lactate secretion rate analysis, lactate production was measured in triplicate with a YSI 2950 Biochemistry Analyzer (Xylem Analytics) against a standard curve and results were normalized to the average of the 0 Gy condition within biological replicates.

### Luminex multiplex immunoassay

Three independent samples of CM collected 7 days post-RT for irradiated and unirradiated control 3T3 fibroblasts were used in a mouse 32-plex Affymetrix kit (performed by Eve Technologies Corporation – Calgary, Alberta, Canada). **Supplementary Table S1** lists all cytokines analyzed, and those with fold change greater than 1.5 after radiation were considered for further analysis.

### Cell lysate collection, SDS-PAGE, and Western blotting

Cell lysates were collected from cells plated in 10 cm dishes, washed twice with ice cold PBS, and incubated with RIPA Buffer (Sigma-Aldrich R0278) containing 5 mM EDTA (Corning 46-034-CI), 1 EDTA-free mini cOmplete EASYpack tablet (Roche 04693159001), and 1 mini PhoSTOP EASYpack tablet (Roche 04906845001). Following mechanical cell disruption in lysis buffer, supernatant was collected and incubated on ice for 30 minutes with constant agitation. Supernatant was sonicated twice (Fisher Scientific Sonic Dismembrator Model 100) and spun down at 4°C at 13,300xg. Supernatant was collected and stored at -80°C until use. A sample from cell lysates was used to determine protein concentration using a bicinchoninic acid assay (Pierce BCA Protein Assay Kit, Thermo Scientific 23227) against a bovine serum albumin standard curve. Samples were denatured in 4x protein loading buffer (LICOR 928-40004) with 10% β-mercaptoethanol and heated at 99°C for 10 minutes. Samples were cooled and loaded into Bio-Rad Mini-PROTEAN TGX pre-cast gels at either 10% (Bio-Rad 4561034) or 4-20% gradient (Bio-Rad 4561094) acrylamide with equivalent total protein across all lanes. A Precision Plus Protein Kaleidoscope Prestained Protein Standard (Bio-Rad 1610375) was additionally loaded to later determine molecular weights of stained samples. Gel electrophoresis was performed using a Bio-Rad PowerPac HC and mini-PROTEAN Tetra system. Samples were run at a constant 200V in a buffer containing TRIS base (Research Products International [RPI] T60040), SDS (RPI L22010), and glycine (RPI G36050). Protein transfer was accomplished using either a 0.2 or 0.45 µm polyvinylidene difluoride membrane following methanol activation and equilibration in transfer buffer containing TRIS base, glycine, and 20% methanol. Transfer occurred either overnight (constant 34V) or within 30 minutes (constant 100V) depending on the protein target as detailed below. Following the transfer, membranes were allowed to dry, re-activated in methanol, and blocked for up to 3 hours in Intercept TBS buffer (LICOR 927-60001). Blocked membranes were incubated in primary antibodies for target protein and loading control diluted with Intercept TBS buffer and 0.2% TWEEN 20 (Sigma-Aldrich P1379) at the following concentrations: anti-LC3B (1:1000, Abcam ab192890, 30 minute transfer), anti-phospho-PINK1 (1:1000, Cell Signaling 89010S, overnight transfer), anti-p62 (1:1000, Cell Signaling Technology 5114, overnight transfer), anti-ATG5 (1:1000, Cell Signaling Technology 12994S, overnight transfer), anti-αTubulin (1:1000, Invitrogen MA1-80017), and anti-GAPDH (1:2000, Biolegend 607902). After overnight incubation at 4°C, samples were washed in TRIS-buffered saline with 0.2% TWEEN-20 three times and then incubated for 1 hour at room temperature in secondary antibody solutions containing: donkey anti-rabbit IRDye 800CW (1:15000, LICOR 926-32213), goat anti-rat IRDye 680RD (1:20000, LICOR 926-68076), or donkey anti-goat IRDye 680LT (1:20000, LICOR 926-68024). Imaging was performed on the Odyssey Fc imager (LICOR) and analysis to determine protein expression relative to loading controls was performed using Image Studio 6.0 (LICORbio).

### Real-time reverse transcription quantitative PCR (RT-qPCR)

Cells were lysed in Trizol (Invitrogen 155960) and RNA was isolated using the PureLink RNA Mini Kit (Invitrogen 12183025) according to the manufacturer’s protocol. For RT-qPCR, 500ng of total RNA was reverse transcribed using High-Capacity cDNA Reverse Transcription Kit (Applied Biosystems 4368814) according to the manufacturer’s protocol. PowerUp™ SYBR™ Green Master Mix (Applied Biosystems A25742) and Custom DNA Oligos (Sigma Aldrich VC00021) were used for gene expression analysis, where genes were normalized to the housekeeping genes *Actb* (murine) or *ACTB* (human) (**Table S2**). RT-qPCR reactions were carried out using a QuantStudio 5 (Applied Biosystems) thermal cycler using the following steps: hold at 95°C for 15 minutes; 40 cycles of alternate denaturation at 95°C for 15 seconds and annealing/extension at 60°C for 1 minute. Melt curve analyses were performed with each experiment to ensure specificity of primers. Relative gene expression was calculated as 2−dCt, where dCt = Ct (test gene) – Ct (housekeeping gene), and Ct is the threshold cycle number. All reactions were performed in triplicate with the average relative expression calculated for each biological replicate.

### TCGA patient data analysis

The TCGA BRCA dataset^82^ was accessed through the University of California Santa Cruz Xena platform on April 1^st^, 2025 to evaluate how gene expression of *GDF15* and *DDIT3* in the normal adjacent tissue (NAT) of breast cancer patients correlated with overall survival^81^. The Xena visualization tool was utilized (https://xenabrowser.net/) to first access the TCGA BRCA dataset, and then only Solid Tissue Normal sample types were kept for analysis. Subsequently, “Breast” was selected as the primary site to only include patient data from TCGA from breast cancer patients. These selections yielded a total patient dataset of n = 113 NAT samples. *GDF15* and *DDIT3* were then entered to evaluate gene expression data via RNAseq analysis, and a Kaplan Meier curve was generated utilizing the built-in functionality of the Xena platform.

### Neutral lipid staining

Oil Red O (Sigma-Aldrich O0625) dye was used for neutral lipid detection in cells. For ORO staining, 0.5 g of ORO powder was added to 100 mL of propylene glycol (Sigma-Aldrich W294004), heated at 97°C for 30 minutes, and stored at room temperature until use. Prior to staining, the ORO solution was vacuum filtered through a 0.45 μm filter (Whatman). Cells on glass coverslips were fixed in neutral-buffered formalin (VWR 16004-128) for 20 minutes at room temperature, washed three times in PBS, and then incubated in propylene glycol for 2 minutes. Following removal of the propylene glycol, coverslips were incubated in ORO solution for 2 hours at room temperature. Samples were then differentiated in 85% propylene glycol for 5 minutes, rinsed in PBS, stained with hematoxylin (Epredia 7211) for 45 seconds, rinsed in DI water, and mounted onto glass slides using a glycerin solution.

### Immunofluorescence staining - cells

For ATP5A1, LAMP-1, and perilipin-2 staining, which required permeabilization, cells on glass coverslips fixed in neutral-buffered formalin were incubated in a 0.1% Triton X-100 solution (Sigma-Aldrich X100) for 10 minutes. For LC3B staining, cells were fixed and permeabilized in 100% ice-cold methanol for 10 minutes at -20°C. Cells were blocked for 1 hour at room temperature with 10% normal goat serum (Vector Labs S-1000) and then incubated overnight at 4°C in a humidified chamber with anti-ATP5A1 (1:500, Proteintech 14676-1-AP), anti-LAMP-1 (1:100, abcam ab208943), anti-perilipin-2 (1:200, Invitrogen MA5-32664), anti-LC3B (1:200, Abcam ab192890) in 1% bovine serum albumin (Sigma A1470) and 0.1% TWEEN 20 (Sigma-Aldrich P1379). Coverslips were mounted onto slides using ProLong Glass Antifade Mountant with NucBlue following secondary antibody incubation with goat anti-rabbit IgG AlexaFluor 488 (1:200 or 1:500, Invitrogen A-11008) or goat anti-rabbit IgG AlexaFluor 594 (1:200 or 1:500, Invitrogen A-11012). A corresponding no primary antibody control was performed for all conditions to confirm specificity.

### Immunofluorescence staining – tissues

For tissue IF staining, tissue sections (5 μm) were deparaffinized and rehydrated followed by antigen retrieval using citrate buffer (10 mM, pH 6). After blocking in 10% normal goat serum, sections were incubated with primary antibodies anti-S100A4 (1:100, R&D Systems MAB4138) and anti-ATP5A1 (1:500) overnight at 4°C. Fluorescently labeled secondary antibodies goat anti-rat IgG AlexaFluor 488 (1:500, abcam ab150157) and goat anti-rabbit AlexaFluor 594 (1:500) were used to stain tissues for 1 hour at room temperature followed by applying a 1X concentration of TrueBlack Lipofuscin Autofluorescence Quencher (Biotium 23007) to reduce autofluorescence. Sections were mounted using ProLong Glass Antifade Mountant with NucBlue and allowed to dry overnight. A corresponding no primary antibody control was performed for all conditions to confirm specificity and determine levels of background tissue autofluorescence.

### Microscopy, image processing, and quantification

Stained samples were imaged using a Leica DMi8 inverted microscope with Leica DFC9000GT sCMOS fluorescence and Leica MC190 HD digital cameras. For fluorescence microscopy, a Lumencor mercury-free SOLA light engine was used for the illumination source. For live cell imaging, cells were imaged using an Okolab Bold Line CO_2_ and temperature unit with air pump. The microscope was fitted with DAPI, GFP, TXR, and Y5 filter cubes. Images (8 bit) were captured using LASX imaging software. Images of perilipin-2, ATP5A1, LAMP-1, LC3B, and S100A4 staining were captured as Z-stacks, where 3D deconvolution using the point-spread function was applied to reduce crosstalk between Z planes in the LASX imaging software (AutoQuant^TM^ Deconvolution algorithm licensed from Media Cybernetics Inc). All images were analyzed using Fiji software^123^. For all IF imaging, area of positive staining was determined by the no primary control images and thresholded accordingly. For perilipin-2 lipid droplet analysis, ATP5A1 total area, and LC3B autophagosome analysis, cross-sectional area of positive staining was calculated per field of view in Fiji and normalized based on cell count. For ATP5A1 aspect ratio analysis, binary images were created from the positive staining threshold, and the aspect ratio was measured in Fiji using the built-in particle analysis function. For ORO neutral lipid quantification, deconvolution of RGB images to evaluate the ORO-stained area was performed using the built-in H&E vector, and positive area was normalized by number of cells. Z-stacks were merged using maximum intensity projections for LC3B and perilipin-2 and based on the Extended Depth of Field function in LASX software for ATP5A1 mitochondrial analysis.

### BODIPY-labeled palmitate uptake experiments and CD36 inhibition

BODIPY FL C_16_ (4,4-Difluoro-5,7-Dimethyl-4-Bora-3a,4a-Diaza-*s*-Indacene-3-Hexadecanoic Acid, Invitrogen D3821) was used to assess the ability of 3T3s to transport fatty acids into the cell from their surroundings. Uptake experiments were performed using modified pre-established protocols^124^. Briefly, fibroblasts were washed in serum free media and incubated in DMEM containing 0.5% fatty acid free bovine serum albumin (Fisher BP9704100) containing 10 μM BODIPY FL C_16_ for 40 minutes at 37°C. Cells were then washed with 0.5% albumin-containing media and incubated with 1.5 µM of Hoechst 33342 (ThermoScientific) for 20 minutes. Cells were washed and imaged live at 37°C and 5% CO_2_ in Hank’s buffered saline solution (HBSS, Corning 21-023-CV). The relative amount of BODIPY FL C_16_ taken up by irradiated and unirradiated control cells was determined by multiplying the area of positive staining by the intensity and normalizing by cell count using Fiji.

### Lysosomal acidification determination and inhibition by CQ

LysoTracker Red DND-99 (Invitrogen L7528) was used to identify acidification of lysosomal structures. Irradiated or unirradiated 3T3s were seeded in tissue culture treated 24 well plates at appropriate seeding densities based on the timepoint after RT and incubated with 50 nM of LysoTracker Red DND-99 and 1.5 µM of Hoechst 33342 solution in media for 30 minutes at 37°C. Following incubation, cells were washed with and imaged live in HBSS at 37°C and 5% CO_2_. Lysosomal acidification was quantified by calculating the product of the area of positive staining and the staining intensity using Fiji and normalizing by the number of cells per field of view. This area was then normalized by dividing by the average of all conditions within each biological replicate. Chloroquine diphosphate salt (CQ, Sigma-Aldrich C6628) was used as an autophagy inhibitor. 5 hours prior to the indicated timepoint, 3T3 cells were incubated with 100 μM of CQ, and iMFs were incubated with 250 μM of CQ in complete media for a maximum of 6 hours^125^. Control cells were treated with equivalent volumes of PBS.

### Tetramethylrhodamine ethyl ester (TMRE) staining for mitochondrial membrane potential

Cells were plated on CELLSTAR μClear™ 96-well, cell culture-treated microplates (Greiner Bio-One 655090) at the appropriate timepoints after RT. Cells were washed with PBS and stained with a 500 nM TMRE (Invitrogen T669) solution in serum-free DMEM for 10 minutes at 37°C and counterstained with Hoechst. Following TMRE incubation, cells were washed twice with PBS and imaged live in PBS using fluorescence microscopy. Localized intensity values were taken at multiple points within individual cells and averaged together across biological replicates.

### MitoTracker Deep Red and MitoSOX staining

MitoTracker Deep Red (Invitrogen M22426) was used to evaluate total mitochondrial activity in 3T3 cells. Stock powder was dissolved in DMSO and used at a working concentration of 200 nM. Cells were washed with PBS and stained with MitoTracker Deep Red in serum free media for 30 minutes at 37°C. Following incubation, cells were washed twice with complete media and then prepared for flow cytometry analysis. MitoSOX™ Red Mitochondrial Superoxide Indicator (Invitrogen M36008) was used to evaluate mitochondrial superoxide production. Stock powder was dissolved in DMSO and used at a working concentration of 5 µM. Cells were washed with PBS and stained with MitoSOX Red in serum free media for 30 minutes at 37°C. Confirmation of successful MitoSOX positive staining was determined by incubation with Antimycin A (Sigma A8674). Following incubation, cells were washed twice with complete media and then prepared for flow cytometry analysis.

### Flow cytometry

Fibroblasts were analyzed via flow cytometry to evaluate MitoTracker Deep Red and MitoSOX staining following RT with or without autophagy inhibition. At either 3 or 7 days after radiation and/or autophagy inhibition, and after staining for the appropriate fluorescent marker, cells were trypsinized, pelleted, and resuspended in ice cold PBS at a concentration of 1x10^6^ cells/mL. Cells were then analyzed live on the Amnis CellStream flow cytometer (Cytek Biosciences). Cells were gated on live and single cells. Flow cytometry data analysis was performed using FlowJo software. Fluorescence intensity was normalized to the mode for each sample.

### Metabolic flux analysis with autophagy and FAO inhibition

Fibroblast mitochondrial oxidative phosphorylation was evaluated through the real-time measurement of the oxygen consumption rate (OCR) and glycolysis through the measurement of the extracellular acidification rate (ECAR) using the Seahorse XFe96 Extracellular Flux Analyzer (Agilent) at the Vanderbilt High Through Screening Core. Fibroblasts irradiated to a dose of 10 Gy (2,200 cells/well) or unirradiated cells (2,200 cells/well) were seeded in an XFe96 culture microplate (Agilent 101085-004) and incubated overnight in complete growth media. The sensor cartridge from the Extracellular Flux Assay Kit (Agilent) was hydrated in sterile DI water overnight at 37°C in a non-CO_2_ incubator. Prior to the experiment, cells being treated were incubated with basal fibroblast media with either 100 μM CQ (6 hours), 40 μM etomoxir sodium salt hydrate (Sigma-Aldrich E1905) for FAO inhibition (24 hours), or an equivalent volume of PBS. The sensor cartridge was incubated with Seahorse XF Calibrant (Agilent 100840-000) 1 hour prior to the experiment, and fibroblasts were washed and then incubated for 1 hour in a non-CO_2_ incubator in XF DMEM Medium, pH 7.4 (Agilent 103575-100) supplemented with 2.0 mM L-glutamine (Sigma-Aldrich G8540), 1.0 mM sodium pyruvate (Sigma-Aldrich S8636), and 10 mM D-(+)-glucose (Sigma-Aldrich G7528). The Seahorse XF Cell Mito Stress Test (Agilent 103015-100) or the Seahorse XF Glycolysis Stress Test (without sodium pyruvate or glucose, Agilent 103020-100) were then performed, and injection strategies were followed per the manufacturer’s instructions with an additional injection step of pure media at the beginning of the assay to confirm no issues with measurement following injection. The following final concentrations of injected drugs and metabolites for both the Mito and Glycolysis Stress Tests were determined from initial optimization experiments: 1.5 μM oligomycin to block ATP synthase, 1.0 μM carbonyl cyanide-p-trifluoromethoxyphenylhydrazone (FCCP) to demonstrate maximal respiration, 0.5 μM of a mixture of rotenone and antimycin A to determine non-mitochondrial respiration, 100 mM glucose for the Glycolytic Stress Test, and 500 mM 2-deoxyglucose to block glycolysis. Cells were counted following the experiment to normalize data per 1,000 cells. Data were analyzed using Seahorse Wave Desktop Software (Agilent), where a correction from a minimum of four background wells were applied. The following calculations were performed on a per well basis: ATP-linked respiration was calculated as the difference between the baseline and post-oligomycin injected OCRs; non-mitochondrial OCR was determined by levels following rotenone and antimycin A injection; and glycolytic ECAR was determined as the difference between the baseline and post-glucose injected ECARs.

### Scratch wound assay

For monoculture, 4T1 cells were plated in 2-well culture inserts in μ-dishes (Ibidi 81176) at 2x10^5^ cells/mL in normal culture media for 36 hours, after which inserts were removed to create a cell-free gap representing the wound scratch. The μ-dishes were washed with PBS and incubated with 7-day post-RT 3T3 CM. Phase contrast images were taken up to 15 hours.

For co-culture, 4T1 cells were labeled with CellTracker^TM^ Deep Red (Invitrogen C34565) and co-cultured with unlabeled 3T3 fibroblasts in a 50/50 mixture in both wells of the μ-dishes starting at 6 days after RT. After 24 hours, inserts were removed to create the cell-free gap. Phase and fluorescence microscopy images were taken after closure, approximately 25 hours. The scratch wound area was calculated in Fiji as the cell-free space between the leading edge of the cells. Scratch wound closure percentage was calculated by dividing scratch wound area by the area at the start of the assay.

### Tumorsphere assays

For monoculture, 4T1 cells were plated at 2x10^4^ cells per well in 200 µL of 7-day post-RT fibroblast CM in a Nunclon Sphera ultra-low attachment 96-well U-bottom plate (Thermo Scientific 174929), allowing the cells to form spheroids. Tumorsphere outgrowth was calculated based on cross-sectional area analyzed from phase contrast images taken 9 days after plating. For co-culture, 1x10^4^ 4T1 cells were plated with 1x10^4^ irradiated or unirradiated shScr or shAtg5 fibroblast cells at 3 days after RT. Co-culture spheroids were grown for 10 days after starting co-culture.

### Statistical analysis

The log-rank test was used to determine statistical significance in Kaplan-Meier analyses. A Mood’s median test for non-equal variances was used to evaluate mitochondrial aspect ratio at 3 and 7 days after RT in unirradiated and irradiated 3T3 fibroblasts. A Mann-Whitney test for non-normally distributed data with similarly shaped distributions was used to analyze 4T1 tumorsphere outgrowth in wild type fibroblast CM. An unpaired, 2-tailed, 2-sample t-test was used within timepoints with an α of 0.05 to evaluate perilipin-2 and ORO lipid droplet analyses without inhibition, BODIPY C16 uptake, 3T3 7-day post-RT baseline ECAR, 3T3 7-day post-RT glycolytic ECAR, the concentration of L-lactate in fibroblast CM, the relative L-lactate secretion rate of irradiated fibroblasts, and the 4T1 scratch-wound closure percentages. A general linear model (one-way ANOVA) with Tukey simultaneous tests for differences of means was used to analyze mitochondrial area per cell, Western blot analysis of phospho-PINK1, ATG5, and p62 protein expressions, LysoTracker staining, 7-day post-RT relative autophagosome area quantification, TMRE measurements of 3-day post-RT wild-type 3T3s, all lipid droplet analyses with CQ incubation, shScr and shAtg5 fibroblast ATP-linked OCR, 7-day ATP-linked OCR with etomoxir, non-mitochondrial OCR, and 4T1 tumorsphere outgrowth in shScr and shAtg5 CM. A one-way ANOVA with Games-Howell pairwise comparison corrections for non-equal variances was used to evaluate 3T3 ATP-linked OCR with CQ incubation, 3-day ATP-linked OCR with etomoxir incubation, 7-day post-RT relative autophagosome area quantification, mitochondrial aspect ratio at 3 and 7 days after RT in CQ treated 3T3 fibroblasts and shAtg5 and shScr cells, and outgrowth of tumorspheres containing 4T1 cells co-cultured with shScr and shAtg5 fibroblasts. A two-way ANOVA with Sidak’s test for multiple comparisons between dose and time or one-way ANOVA with Sidak’s test for multiple comparisons for comparison between dose was used to analyze gene expression data. All analyses were performed using Minitab Statistical Software Version 22 and GraphPad Prism 10.

## Resource Availability

**Lead contact.** Further information and requests for resources and reagents should be directed to and will be fulfilled by the Lead Contact, Marjan Rafat.

**Materials availability.** This study did not generate new unique reagents.

**Data and code availability**. This study did not generate any unique datasets or code.

