## Supplementary Figures for "Radiation-Induced Autophagy Regulates the Fibroblast Mitochondrial Stress Response and Crosstalk with Triple-Negative Breast Cancer Cells"

### Supplemental Figures and Tables

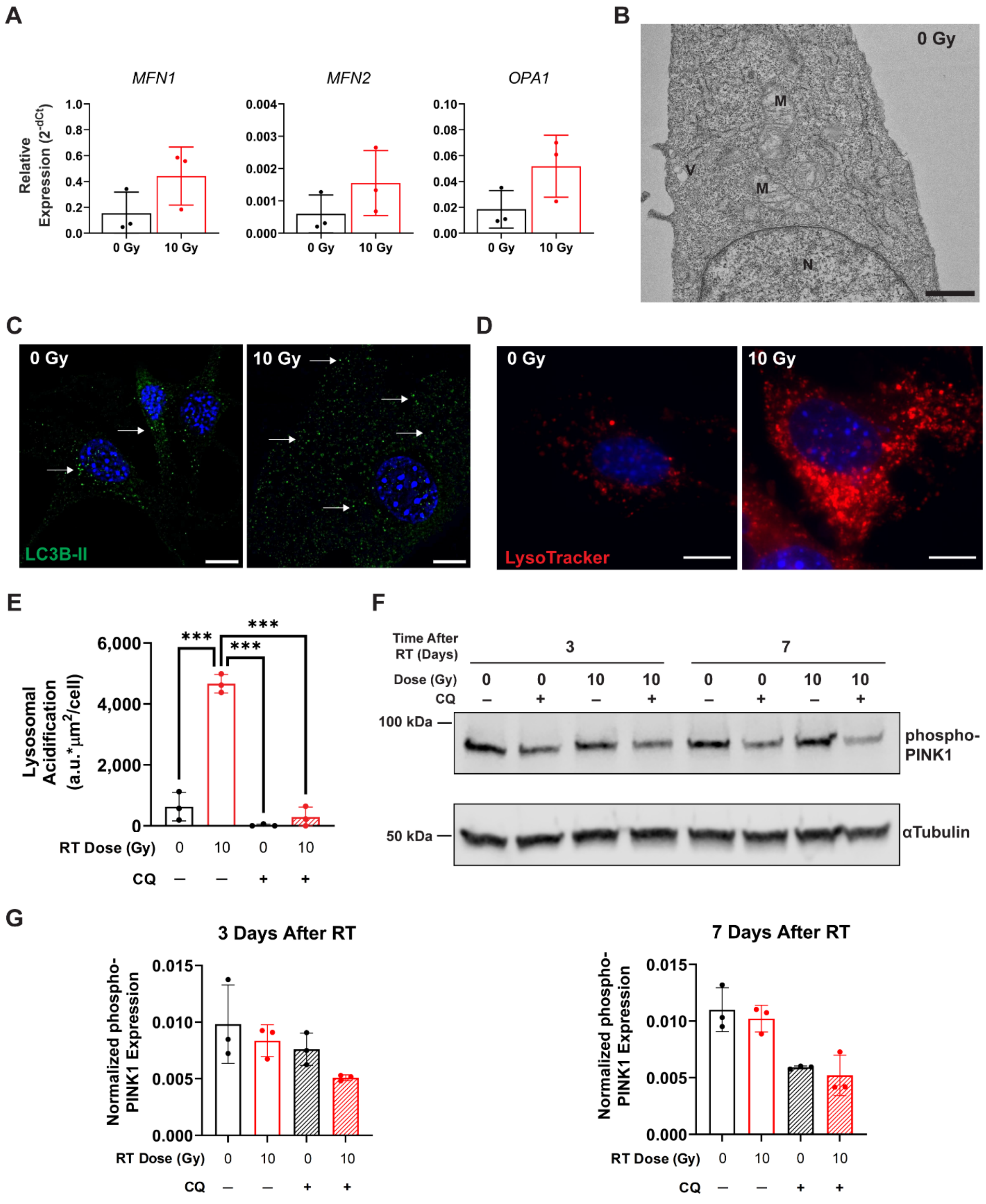

**Figure S1. Autophagy and mitochondrial elongation are linked in irradiated fibroblasts. (A)** Relative gene expression of fusion-related genes *MFN1*, *MFN2*, and *OPA1* in unirradiated (0 Gy) and irradiated (10 Gy) human immortalized mammary fibroblasts (iMFs) 7 days after radiation therapy (RT) (n=3). **(B)** Representative transmission electron microscopy (TEM) image of unirradiated 3T3 fibroblasts 7 days after RT. N=nucleus, M=mitochondria, V=vacuole-like structure. Scale bar is 800 nm. **(C)** Representative images of LC3B (green) and nuclear (blue) staining in irradiated and unirradiated 3T3s 7 days after RT. Autophagosomes are identified by the formation of bright LC3B puncta (white arrows). Scale bars are 10 µm. **(D)** Representative images of LysoTracker Red DND-99 (red) and nuclear (blue) live cell staining in irradiated and control 3T3s 3 days after RT. Scale bars are 10 µm. **(E)** Quantification of live cell lysosomal staining in irradiated and control 3T3 fibroblasts with and without chloroquine (CQ) incubation (n=3). **(F)** Representative Western blot showing phosphorylated PINK1 expression with αTubulin loading control in 3T3 fibroblasts 3 and 7 days after RT with and without CQ incubation. **(G)** Quantification of phosphorylated PINK1 expression normalized to αTubulin in 3T3 fibroblasts 3 and 7 days after RT with and without CQ incubation (n=3). Statistical analysis in **(E)** was determined by one-way ANOVA and Tukey simultaneous tests for differences of means where ***$p<0.001$. Error bars show standard deviation.

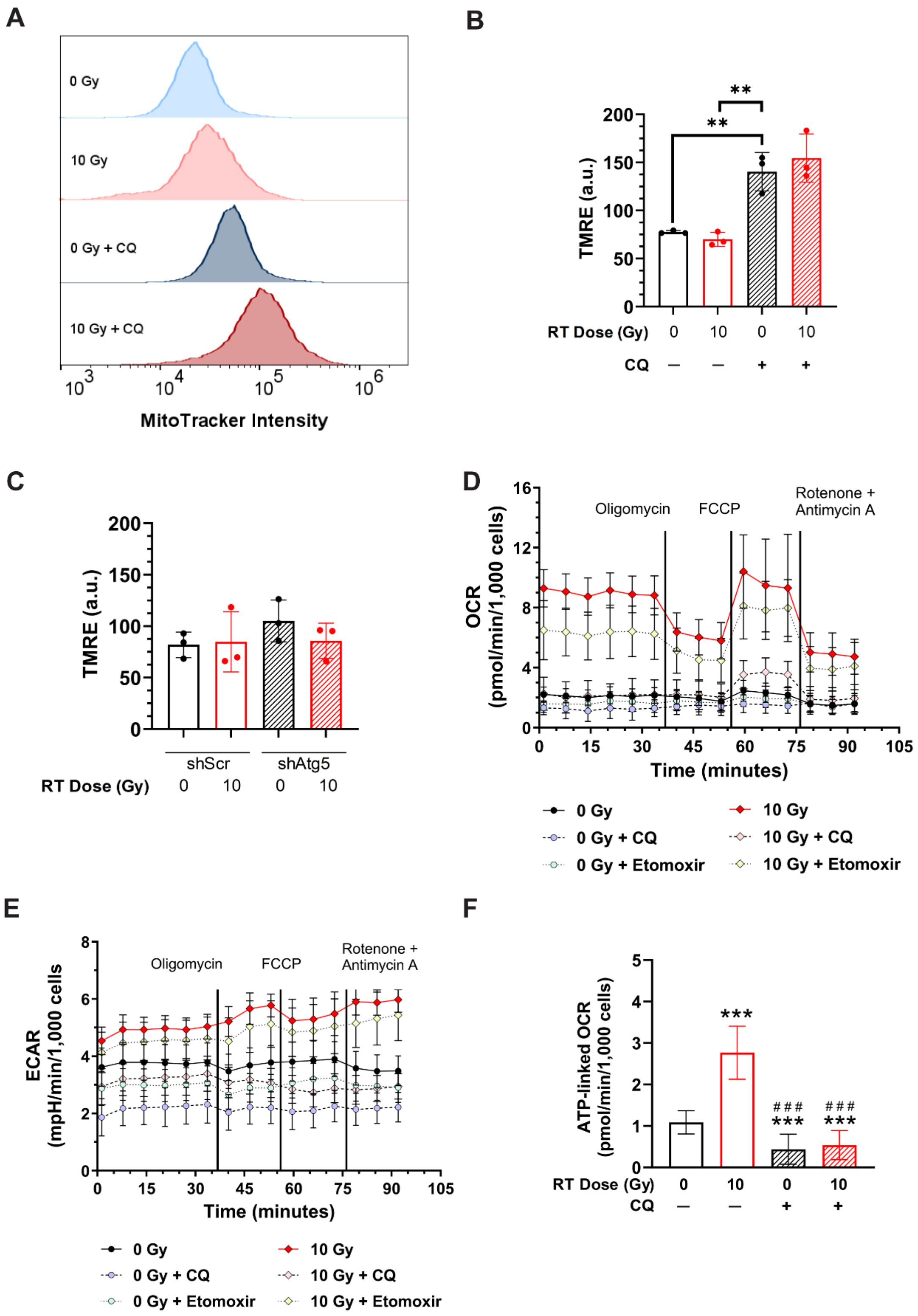


**Figure S2. Evaluating irradiated fibroblast bioenergetics. (A)** Representative flow cytometry analysis of irradiated and unirradiated 3T3 fibroblasts stained with MitoTracker Deep Red at 3 days after RT following incubation with CQ. **(B–C)** Tetramethylrhodamine ethyl ester **(**TMRE**)** intensity measurements of 3T3 fibroblasts **(B)** with and without CQ incubation (n=3) and **(C)** in shScr and shAtg5 fibroblasts 3 days after RT (n=3). Representative OCR **(D)** and ECAR **(E)** curves of irradiated and unirradiated 3T3 fibroblasts with and without CQ or etomoxir incubation to inhibit autophagy or fatty acid oxidation, respectively, from a modified mitochondrial stress test. **(F)** ATP-linked oxygen consumption rate (OCR) in irradiated and unirradiated 3T3s with and without CQ incubation 3 days after RT (n=3). Statistical analysis for **B** determined by one-way ANOVA with Tukey simultaneous tests for differences of means with **$p<0.01$. Statistical analysis for **F** determined by one-way ANOVA with Games-Howell pairwise comparison corrections for non-equal variances with ***$p<0.001$. Error bars are standard deviation.

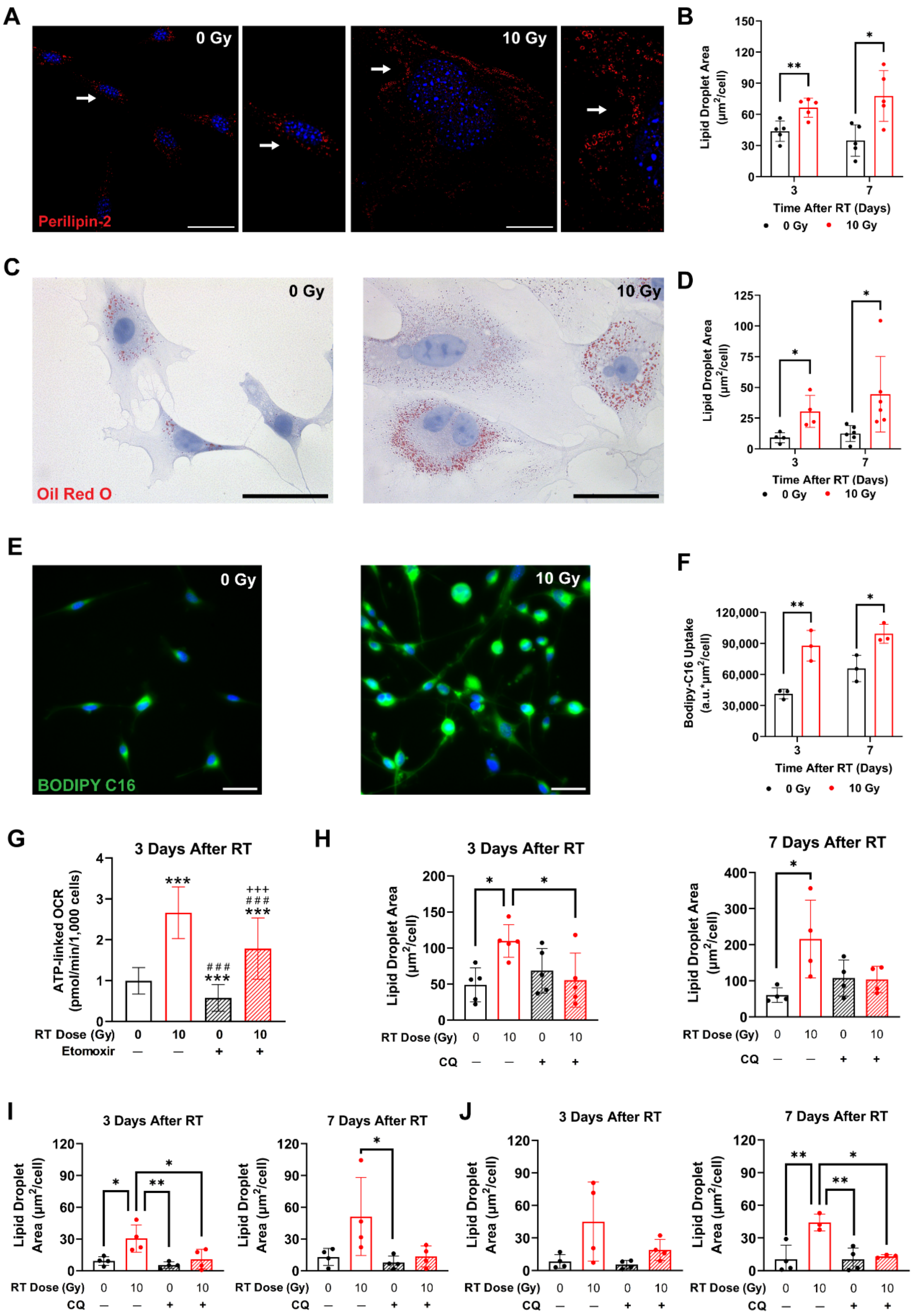


**Figure S3. Autophagy and fatty acids regulate irradiated fibroblast bioenergetics. (A)** Representative images of perilipin-2 (red) and nuclear (blue) staining in 7-day post-RT fibroblasts. White arrows represent areas of increased magnification. Scale bars are 30 µm. **(B)** Quantification of the cross-sectional area of lipid droplets from perilipin-2 IF staining of 3T3s (n=5). **(C)** Representative images of 3T3s stained with Oil Red O (ORO) and counterstained with hematoxylin at 7 days after RT. **(D)** Quantification of the lipid droplet area positively stained with ORO in 3T3s and normalized per cell (n=4, 3 day; n=6, 7 day). **(E)** Representative images of 3T3s 3 days after RT following BODIPY C16 (green) uptake with nuclear staining (blue). Scale bars are 30 µm. **(F)** Quantification of fluorescence images of BODIPY-labeled palmitate uptake in irradiated and control 3T3s up to 7 days after RT (n=3). **(G)** Quantification of the ATP-linked OCR following incubation with etomoxir to determine FAO in 3T3 fibroblast 3 days after RT. **(H)** Quantification of the cross-sectional lipid droplet area determined from IF staining of perilipin-2 in 3T3 fibroblasts at 3 and 7 days after RT with CQ incubation to inhibit autophagy (n=5). **(I)** Quantification of the lipid droplet area positively stained with ORO in irradiated and control 3T3s at 3 and 7 days after RT with and without CQ incubation (n=4). **(J)** Quantification of ORO staining in irradiated and control iMFs at 3 and 7 days after RT with and without CQ incubation (n=3–4). For **B**, **D**, and **F**, statistical significance was determined by an unpaired two-tailed t-test within timepoints with *p<0.05 and **p<0.01. For **G**, statistical significance was determined by one-way ANOVA with Games-Howell pairwise comparison corrections for non-equal variances. Symbols for pairwise comparisons: * for comparison to 0 Gy & 0 µM etomoxir, # for comparison to 10 Gy & 0 µM etomoxir, and + for comparison to 0 Gy & 40 µM etomoxir, where $^{***, \#\#\#, +++}$p<0.001. For **H–J**, statistical analysis was determined by one-way ANOVA and Tukey simultaneous tests for differences of means within each timepoint, where *p<0.05 and **p<0.001. Error bars show standard deviation.

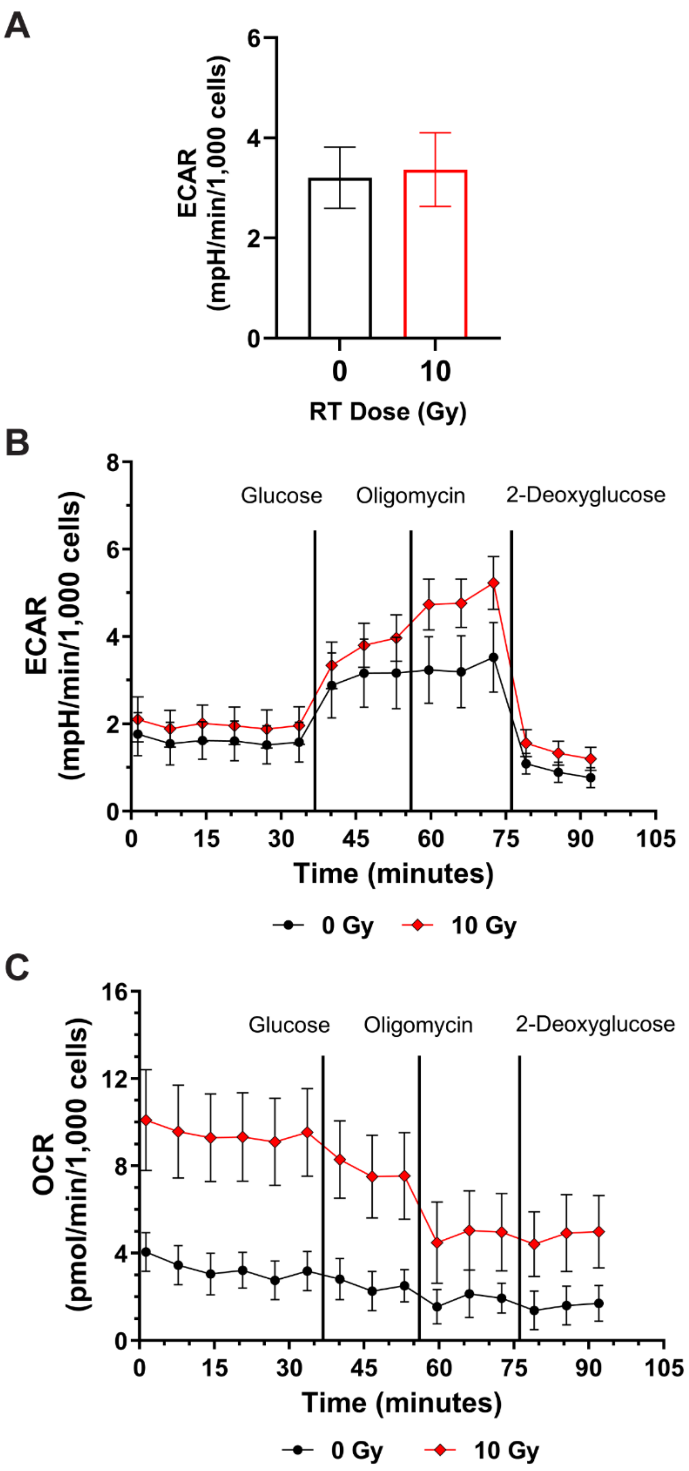


**Figure. S4. Irradiated fibroblasts have increased extracellular acidification rate (ECAR) at late timepoints after RT. (A)** Baseline ECAR in 3T3 fibroblasts from a mitochondrial stress test at 3 days after RT (n=3). Representative ECAR **(B)** and OCR **(C)** curves of irradiated and unirradiated 3T3 fibroblasts with and without CQ incubation to inhibit autophagy from a modified glycolytic stress test. Error bars are standard deviation.

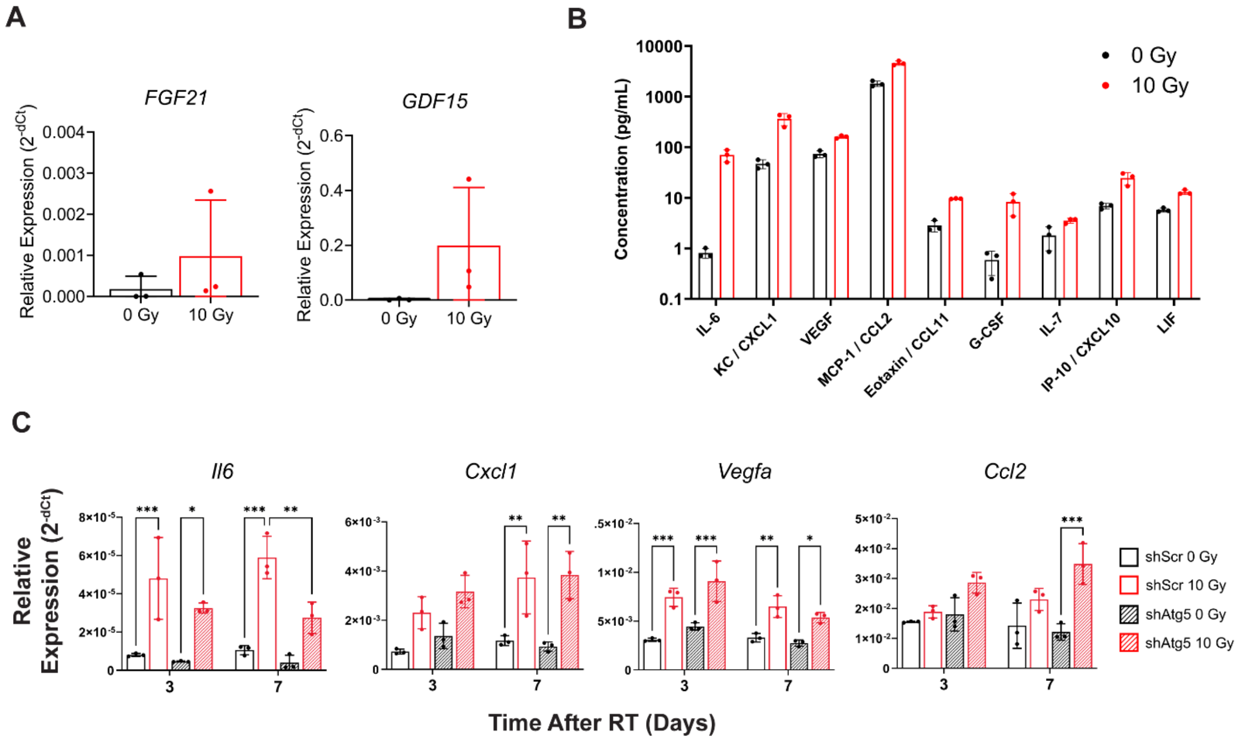


**Figure S5. Autophagy promotes differential expression of mitokines, cytokines, and chemokines in irradiated fibroblasts. (A)** Relative gene expression of *FGF21* and *GDF15* in human iMFs 7 days after RT. **(B)** Multiplex analysis of upregulated cytokines and chemokines in CM from irradiated and unirradiated 3T3 fibroblasts 7 days after RT (n=3). See **Table S1** for evaluated cytokines. **(C)** Relative gene expression of *Il6*, *Cxcl1*, *Vegfa*, and *Ccl2* at 3 and 7 days after RT in shScr and shAtg5 fibroblasts (n=3). Statistical analysis in **C** determined by a 2-way ANOVA with multiple comparisons by the Sidak method with *p<0.05, **p<0.01, and ***p<0.001. Error bars are standard deviation.

**Table S1. List of cytokines evaluated using a cytokine 32-plex immunoassay.**

| **Fold Change > 1.5** | |
|---|---|
| **Name** | **Abbreviation** |
| Eotaxin | Eotaxin |
| Granulocyte colony-stimulating factor | G-CSF |
| Monocyte chemoattractant protein-1/<br>C-C Motif chemokine ligand 2 | MCP-1/<br>CCL2 |
| Interleukin 6 | IL-6 |
| Leukemia inhibitory factor | LIF |
| Interferon γ-induced protein 10 kDa/<br>Chemokine (C-X-C motif) ligand 10 | IP-10/<br>CXCL10 |
| Interleukin 7 | IL-7 |
| Interleukin 15 | IL-15 |
| Keratinocytes-derived chemokine | KC |
| Vascular Endothelial Growth Factor | VEGF |
| **Fold Change < 1.5 or Out Of Range** | |
| **Name** | **Abbreviation** |
| Interferon gamma | IFNγ |
| Interleukin 1 beta | IL-1β |
| Interleukin 2 | IL-2 |
| Interleukin 3 | IL-3 |
| interleukin 4 | IL-4 |
| Interleukin 5 | IL-5 |
| Interleukin 9 | IL-9 |
| Interleukin 10 | IL-10 |
| Interleukin 12p40 | IL-12p40 |
| Interleukin 12p70 | IL-12p70 |
| Interleukin 13 | IL-13 |
| Interleukin 17 | IL-17 |
| Macrophage colony-stimulating factor | M-CSF |
| Macrophage inflammatory protein-1 alpha | MIP-1α |
| Macrophage inflammatory protein-1 beta | MIP-1β |
| Macrophage inflammatory protein-2 | MIP-2 |
| Granulocyte-macrophage colony-stimulating factor | GM-CSF |
| Lipopolysaccharide-induced CXC chemokine/<br>Chemokine (C-X-C Motif) ligand 5 | LIX/<br>CXCL5 |
| Monokine induced by gamma interferon/<br>Chemokine (C-X-C Motif) ligand 9 | MIG/<br>CXCL9 |
| Tumor necrosis factor alpha | TNFα |
| Interleukin 1 alpha | IL-1α |
| Regulated on activation, normal T cell expressed and secreted/<br>C-C Motif Chemokine Ligand 5 | RANTES/<br>CCL5 |

**Table S2. qPCR primer sequences.**

| Gene | Forward Primer | Reverse Primer |
|---|---|---|
| *ACTB* | GCCTCGCCTTTGCCGAT | AGGTAGTCAGTCAGGTCCCG |
| *Actb* | CCACCATGTACCCAGGCATT | CGGACTCATCGTACTCCTGC |
| *Atf5* | CCAATTGTTGGTGCAGCCTC | CTTCTTTTGCTTGCGGTCCC |
| *Ccl2* | CAGGTCCCTGTCATGCTTCT | GTGGGGCGTTAACTGCATCT |
| *Ddit3* | CCACCACACCTGAAAGCAGAA | AGGTGAAAGGCAGGGACTCA |
| *Cxcl1* | ACTCAAGAATGGTCGCGAGG | GTGCCATCAGAGCAGTCTGT |
| *FGF21* | CAGGTGAGAGGCTTCCAAGG | TGCGCCCCATCTGAATTTCT |
| *Fgf21* | CCTCCAGTTTGGGGGTCAAG | ACCACTGTTCCATCCTCCCT |
| *GDF15* | GCAAGAACTCAGGACGGTGA | TGGAGTCTTCGGAGTGCAAC |
| *Gdf15* | AAAGACACACTCAGGACACAA | GCTTCAGGGGCCTAGTGATG |
| *Hspd1* | CGGAAATGACGCCACTTGAC | CATCCCCAGCCTCTTCGTTT |
| *Il6* | GCCTTCTTGGGACTGATGCT | TGTGACTCCAGCTTATCTCTTGG |
| *MFN1* | GGGTGATAGTTGGAGCGGAG | GAGCTCTTCCCACTGCTTGT |
| *Mfn1* | CCTCTTTCGGGAGGATGAGA | AGGGCACATTTTGCTTTGGG |
| *MFN2* | GGAAGGTGAAGCGCAATGT | AGAGTTGGGCCACATCACAC |
| *Mfn2* | TCTCCATTCAAGAAGCTTGGACA | GCTGATACCCCTGACCTTGG |
| *OPA1* | GGCTCTTGCGGAAGTCCAT | TGCTGGCCAAAAATTCCTGC |
| *Opa1* | TTCTGAGGCCCTTCTCTTGT | ATGCCTGATGTCACGGTGTT |
| *Sirt5* | GCCACCGACAGATTCAGGTT | CCACAGGGCGGTTAAGAAGT |
| *Vegfa* | AAAACACAGACTCGCGTTGC | GGTCTTTCCGGTGAGAGGTC |
